# Systematic reconstruction of the cellular trajectories of mammalian embryogenesis

**DOI:** 10.1101/2021.06.08.447626

**Authors:** Chengxiang Qiu, Junyue Cao, Tony Li, Sanjay Srivatsan, Xingfan Huang, Diego Calderon, William Stafford Noble, Christine M. Disteche, Malte Spielmann, Cecilia B. Moens, Cole Trapnell, Jay Shendure

## Abstract

Mammalian embryogenesis is characterized by rapid cellular proliferation and diversification. Within a few weeks, a single cell zygote gives rise to millions of cells expressing a panoply of molecular programs, including much of the diversity that will subsequently be present in adult tissues. Although intensively studied, a comprehensive delineation of the major cellular trajectories that comprise mammalian development *in vivo* remains elusive. Here we set out to integrate several single cell RNA-seq datasets (scRNA-seq) that collectively span mouse gastrulation and organogenesis. We define cell states at each of 19 successive stages spanning E3.5 to E13.5, heuristically connect them with their pseudo-ancestors and pseudo-descendants, and for a subset of stages, deconvolve their approximate spatial distributions. Despite being constructed through automated procedures, the resulting trajectories <u>o</u>f <u>m</u>ammalian <u>e</u>mbryogenesis (TOME) are largely consistent with our contemporary understanding of mammalian development. We leverage TOME to nominate transcription factors (TF) and TF motifs as key regulators of each branch point at which a new cell type emerges. Finally, to facilitate comparisons across vertebrates, we apply the same procedures to single cell datasets of zebrafish and frog embryogenesis, and nominate “cell type homologs” based on shared regulators and transcriptional states.

## Introduction

A fundamental goal of developmental biology is to understand the relationships of cell types to one another during embryogenesis, as well as the molecular programs that underlie each cell type’s emergence. In principle, developmental programs can be comprehensively described, *e.g.* Sulston and colleagues’ reconstruction of the complete embryonic lineage of the roundworm *C. elegans* through visual observation (Sulston et al. 1983). However, *C. elegans* -- small, translucent, and developmentally invariant -- remains the only model organism for which such a complete description has been realized.

Over the past four years, we and others have developed and applied new technologies for single cell molecular profiling to developing model organisms at the “whole animal” scale, including the worm, fly, zebrafish, frog, and mouse (Pijuan-Sala et al. 2019; Cao et al. 2019; Wagner et al. 2018; Briggs et al. 2018; Farrell et al. 2018; Cao et al. 2017). Such studies lay the foundations for global views of metazoan development, including, for example, populating the Sulston lineage with the gene expression programs of each cell type (Cao et al. 2017; Packer et al. 2019).

For mouse embryogenesis in particular, we and others have performed single cell or single nucleus RNA-seq data (scRNA-seq) during implantation (Cheng et al. 2019; Mohammed et al. 2017), gastrulation (Pijuan-Sala et al. 2019) and organogenesis (Cao et al. 2019). Collectively, these four studies span the development of the mouse embryo from dozens of cells of a few types (E3.5) to millions of cells of hundreds of types (E13.5). However, the data associated with these studies has yet to be systematically integrated in a manner that permits their robust exploration. Such integration is challenging, both for technical reasons (*e.g.* different studies, different technologies, batch effects, etc.) as well as because of the sheer complexity of mouse development.

Here we set out to systematically reconstruct the major cellular trajectories of mammalian embryogenesis from E3.5 to E13.5. Our primary strategy is inspired by Briggs and colleagues (Briggs et al. 2018) and makes several assumptions: 1) Although mouse development is variable, key patterns will be invariant across wild-type animals; 2) “*Omnis cellula e cellula*” also applies to cell states, *i.e.* cell states observed at a given timepoint must have arisen from cell states present at the preceding timepoint; 3) We are sampling frequently and deeply enough that newly detected cell states will not arise from antecedent cell states that were undetected at the preceding timepoint; 4) Provided that the delta in time is small enough, transcriptional similarity is an effective means of linking related cell states observed at adjacent timepoints.

A caution is that in contrast to the Sulston’s seminal map of *C. elegans*, we focus here on reconstructing cellular *trajectories* (Trapnell et al. 2014), a concept related, but by no means equivalent, to cell lineage. Although it is a reasonable expectation that closely related cells (*e.g.* siblings) will be transcriptionally similar (Packer et al. 2019), the converse is not necessarily true. For example, molecular states can be insufficiently divergent, or even convergent, both of which obscure lineage relationships (Wagner and Klein 2020). Furthermore, even the expectation that lineally closely related cells will be transcriptionally similar is not always met, as rapidly changing molecular states can lead to “gaps” in trajectories (Packer et al. 2019). In sum, our goal here is a continuous, navigable roadmap of the molecular states of cell types during mouse development. Such a roadmap may constrain the potential lineage relationships amongst constituent cell types, but it does not explicitly specify them.

### Systematic reconstruction of the cellular trajectories of mouse embryogenesis

The datasets used here are derived from 468 samples (where each sample is an individual embryo or small pool of mouse embryos) from 19 timepoints or stages spanning E3.5 to E13.5, with successive stages separated by as few as 6 hours but no more than 1 day (**Supplementary Table 1**). The number of profiled cells totalled 1,443,099, and ranged from 99 to 449,621 per stage (**Supplementary Fig. 1a-b**). For each stage, we performed pre-processing followed by Louvain clustering and manual annotation of individual clusters based on marker gene expression (**Supplementary Table 2**). Here we use “cell state” to mean an annotated cluster at a given stage. Altogether, we identified 413 cell states across the 19 timepoints, each of which received one of 84 cell type annotations.

For each pair of adjacent stages, we projected cells into a shared embedding space (Stuart et al. 2019). Relative to (Briggs et al. 2018), a technical challenge here is that both within and between some stages, data was generated by different groups using different scRNA-seq technologies. To address this, we performed anchor-based batch correction prior to integration, which proved quite effective, including across platforms as well as across cells vs. nuclei (**Supplementary Fig. 1c-e**) (Stuart et al. 2019). After co-embedding, we applied a *k*-nearest neighbor (*k*-NN) based heuristic to connect cell states between adjacent stages. Briefly, for each cell state at the later timepoint, we identified the 5 closest cells from the antecedent timepoint in the co-embedding. Bootstrapping to obtain a robust estimate (500 iterations with 80% subsampling), we then calculated the median proportion of such neighbors derived from each potential antecedent cell state, and treated this as the weight of the corresponding edge.

As a simple example, clustering and annotation of scRNA-seq data from two adjacent timepoints, E6.25 and E6.5, identified 5 and 6 cell states, respectively (**Fig. 1a**). If we co-embed these data and follow the aforedescribed procedure, we strongly link 5 cell states at E6.5 to 5 cell states bearing the same annotations at E6.25. The new cell state at E6.5, which corresponds to the primitive streak, is strongly linked to E6.25 epiblast, which we assign as its pseudo-ancestor (**Fig. 1a**). Upon applying this procedure to E6.5 → E6.75 and E6.75 → E7.0, the primitive streak is further assigned as the pseudo-ancestor of the nascent mesoderm, anterior primitive streak and primordial germ cells (**Supplementary Fig. 2**).

**Figure 1.**
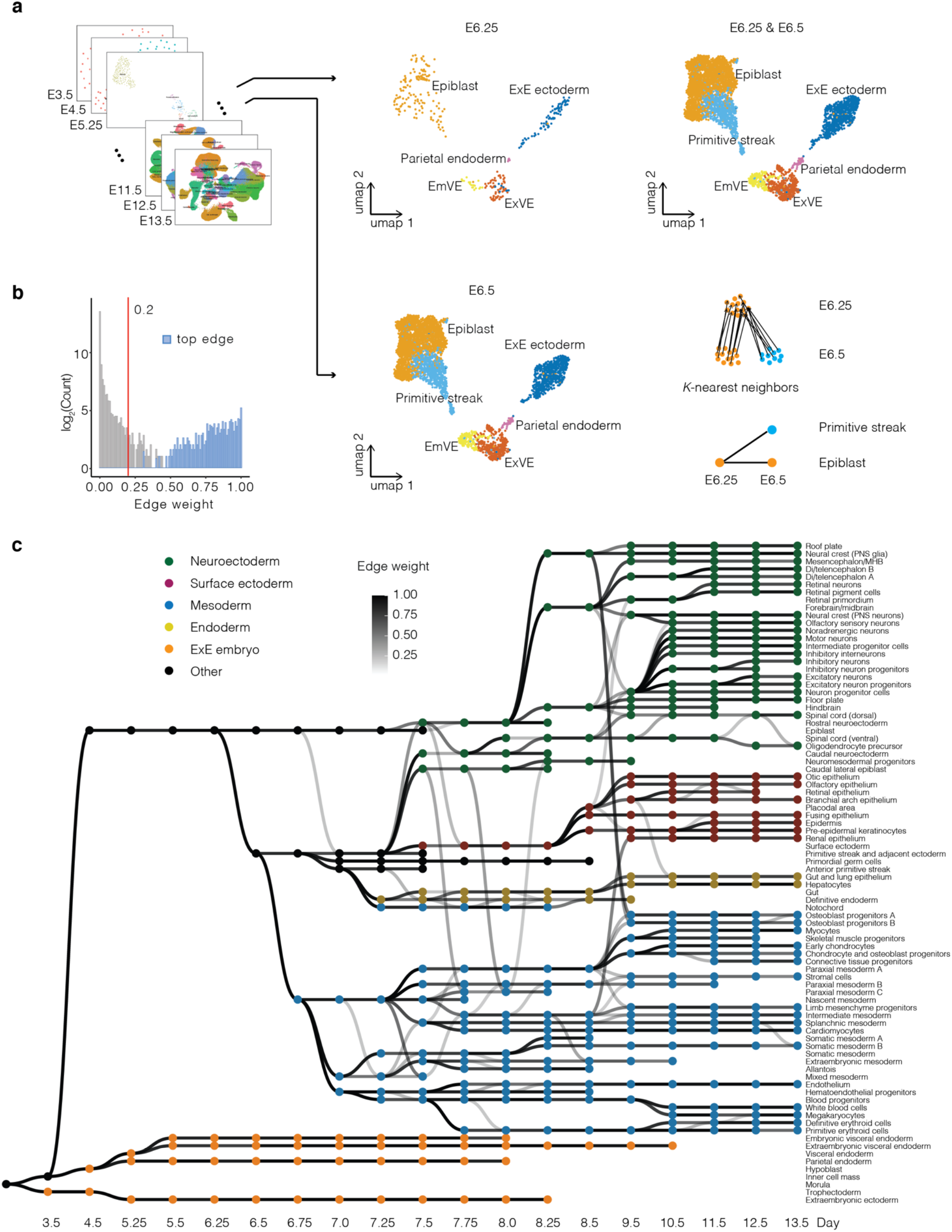
Systematic reconstruction of the cellular trajectories of mouse embryogenesis. **a,** Overview of approach. Cells from each pair of adjacent stages were projected into the same embedding space (Stuart et al. 2019). UMAP visualizations of co-embedded cells from E6.25 and E6.5 are shown separately (middle column) or together (top right). A *k*-NN heuristic was applied to infer one or several pseudo-ancestors for each of the cell states observed at the later time point (bottom right). ExE: extraembryonic. EmVE: embryonic visceral endoderm. ExVE: extraembryonic visceral endoderm. **b,** Histogram of all calculated edge weights. The *y-*axis is on a log2 scale. Edges with weights above 0.2 (red line) were retained. “Top edges” are those with the highest weight amongst all potential antecedents of each cell state. **c,** Directed acyclic graph showing inferred relationships between cell states across early mouse development. Each row corresponds to one of 84 cell type annotations, columns to developmental stages spanning E3.5 to E13.5, nodes to cell states, and node colors to germ layers. All edges with weights above 0.2 are shown in grey scale. Of note, placental tissues were not actively retained during the isolation of embryos from later timepoints (Cao et al. 2019). ExE: extraembryonic. PNS: peripheral nervous system. MHB: midbrain-hindbrain boundary.

We applied this approach across each of the 18 pairs of adjacent timepoints (**Supplementary Fig. 3**). Although the resultant edge weights were bimodally distributed, a cutoff of 0.2 was selected towards being more inclusive of weaker relationships as well as to ensure connectivity of the overall graph (**Fig. 1b**). Of note, we introduced 4 “dummy nodes”, corresponding to the morula at E3.0 (as a root for the trophectoderm and inner cell mass), trophectoderm at E3.5 and E4.5 (which had been removed at these timepoints by immunosurgery (Mohammed et al. 2017)) and parietal endoderm at E6.75 (undetected, likely due to undersampling). The resulting representation is a directed acyclic graph with 417 nodes and 468 edges that captures trajectories <u>o</u>f <u>m</u>ammalian <u>e</u>mbryogenesis (TOME) (**Fig 1c**).

### Do molecular trajectories recapitulate cellular phylogenies?

To reiterate, the graph shown in **Fig. 1c** (TOME) does not reflect cell lineage but rather relationships between cell states that were inferred on the basis of transcriptional similarity. Nonetheless, under the supposition that lineally related cell states diverge from one another through a succession of continuous molecular states, we can ask whether or not established lineage relationships are respected by TOME. Of the 468 edges with weights greater than 0.2, 328 (70%) are between cell states bearing the same annotation, while 140 (30%) are between cell states bearing different annotations. In **Supplementary Table 3**, we show all edge weights and comment on inferred transitions. Several observations merit emphasis.

First, the graph largely respects germ layers, which are indicated by node colors in **Fig. 1c**. There are no edges between extraembryonic and embryonic cell states, and relatively few edges between embryonic cell states of different germ layers. Among the strongest edges that cross between germ layers “boundaries” are two edges that connect E8.5 neural crest (PNS glia) to two subtypes of E9.5 osteoblast progenitors, presumably corresponding to the well-established neural crest contribution to bones (Tani et al. 2020); another edge between E8.5 intermediate mesoderm and E9.5 renal epithelium, also an established contribution across germ layers (Bouchard 2004); and another edge between caudal lateral epiblast and a subset of paraxial mesoderm at E7.5-E8.0, also previously described (Albors, Halley, and Storey 2018)

Second, 80% of cell types are strongly linked (edge weight greater than 0.7) to a single pseudo-ancestor when they first appear. These strong edges generally respect established lineage relationships, *e.g.* parietal and visceral endoderm arising from hypoblast (Rivera-Pérez and Hadjantonakis 2014), notochord and definitive endoderm arising from the anterior primitive streak (Balmer, Nowotschin, and Hadjantonakis 2016; Wells and Melton 1999), cardiomyocytes arising from splanchnic mesoderm (Ivanovitch, Temiño, and Torres 2017), and many others.

Third, apparent convergences — instances wherein we assign more than one pseudo-ancestor to a cell state — sometimes correspond to a given cell type persisting and “contributing” to another cell type over several consecutive timepoints (*e.g.* hemoendothelial progenitors are recurrently assigned as pseudo-ancestors of endothelial cells at E7.75-E8.25). In other cases, apparent convergences may reflect incomplete separation between highly related cell types, rather than ongoing differentiation (*e.g.* the several edges between notochord and definitive endoderm; recurring edges between different subtypes of mesoderm). However, yet other cases reflect bonafide convergence of transcriptional states, *i.e.* where a cell type has multiple origins. For example and as also noted above, the two subtypes of E9.5 osteoblast progenitors have edges back to both E8.5 neural crest and E8.5 paraxial mesoderm, consistent with the literature (Tani et al. 2020), while a subtype of paraxial mesoderm has edges back to nascent mesoderm and caudal lateral epiblast (Albors, Halley, and Storey 2018). Of note, not all established convergences are captured, *e.g.* the known contribution of embryonic visceral endoderm to the gut at E7.5-E7.75 (Nowotschin and Hadjantonakis 2020) is detected at E7.5-E7.75, but falls just short of the 0.2 edge weight threshold (**Supplementary Table 3**).

Fourth, an important limitation of this heuristic approach, made apparent by a few clear inaccuracies in the graph, is that true lineage relationships for a given cell state can be obscured by the presence of a highly similar cell state at the preceding timepoint. For example, E9.5 neuron progenitor cells are assigned as the pseudo-ancestor of multiple neuronal subtypes that appear at E10.5, but we do not observe these same relationships to recur at subsequent timepoints, although neuronal differentiation is surely ongoing. This is probably because at timepoints subsequent to E10.5, each derivative neuronal subtype is most similar to itself at the preceding timepoint, such that it fails to be linked back to the persisting neuron progenitors. This same phenomenon probably explains another error, wherein when definitive erythroid cells first appear at E10.5, they are linked to E9.5 primitive erythroid cells, rather than to blood progenitors. For a more exhaustive consideration of the ways in which trajectory-based inference can be misleading about cell lineage histories, see (Wagner and Klein 2020). Of note, at least some of the inaccuracies noted above are resolvable by focused analyses that leverage the distinction between nascent and spliced transcripts, *i.e.* RNA velocity (La Manno et al. 2018). For example, if we reanalyze these problematic subsets of TOME with *scVelo* (Bergen et al. 2020), the heterogeneity and ongoing contributions of neuron progenitors is much more evident (Sagner et al., n.d.) (**Fig. 2a**), and primitive and definitive hematopoiesis are much more clearly separated (**Fig. 2b**). A third inaccuracy, also potentially explained by the same phenomenon, is lack of a relationship between neural crest neurons and glia. However, once again, if we reanalyze these subsets on their own, their shared origin is much more apparent, as are the multiple waves of sensory neurogenesis (Pavan and Raible 2012) (**Fig. 2c**).

**Figure 2.**
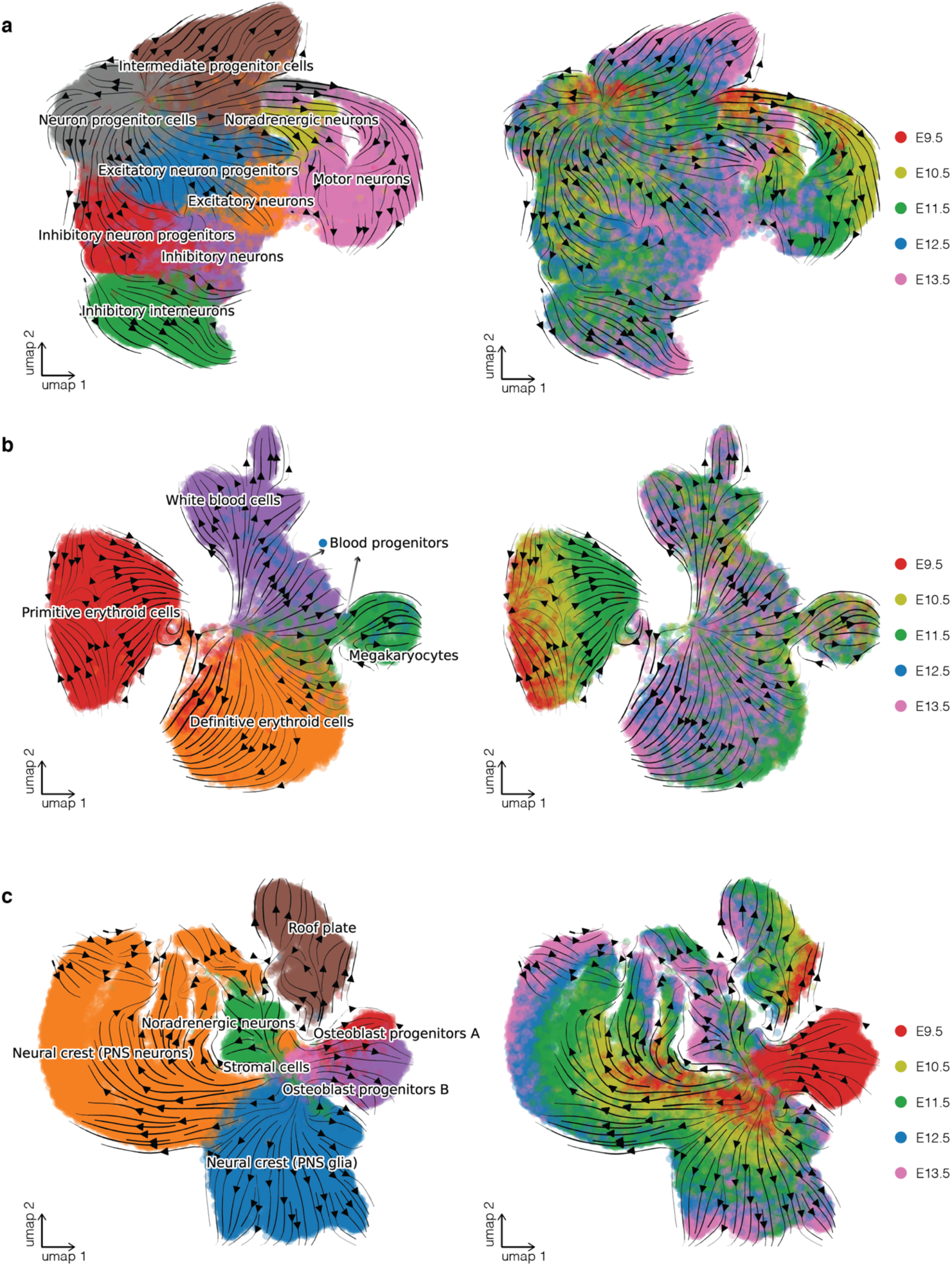
RNA velocity clarifies relationships between cell types during neuronal differentiation, hematopoiesis and neural crest development. **a**, RNA velocity was estimated on the basis of the proportion of reads mapping to exonic vs. intronic portions of genes using *scVelo* (Bergen et al. 2020). Cells corresponding to noradrenergic neurons, motor neurons, intermediate progenitor cells, inhibitory interneurons, inhibitory neuron progenitors, inhibitory neurons, excitatory neuron progenitors, excitatory neurons, and neuron progenitor cells from E9.5 and E13.5 were included in this analysis, after downsampling each cell state to 5,000 cells. UMAP visualization of co-embedded cells and cell state transition trends (arrows) are shown. Smaller panels show the same UMAP visualization but with coloring of cells from individual timepoints. **b**, Same as panel a, but for cells corresponding to blood progenitors, white blood cells, megakaryocytes, definitive erythroid cells and primitive erythroid cells from E9.5 and E13.5. **c**, Same as panel a, but for cells corresponding to neural crest (PNS glia), neural crest (PNS neurons), roof plate, noradrenergic neurons from E9.5-E13.5, as well as stromal cells and osteoblast progenitors A & B from E9.5 only.

Fifth, a further limitation is that our reliance on discrete entities, *i.e.* cell states, obscures aspects of developmental biology that are inherently continuous. For example, spatial transcriptional heterogeneity, which often manifests as continuous gradients, is obscured by cell type or cell state discretization. Here, we have represented aspects of spatial heterogeneity in a limited way through distinct nodes (*e.g.* fore/mid/hindbrain; paraxial mesoderm A/B/C), but this is far from ideal.

In summary, molecular trajectories often recapitulate well-documented cellular phylogenies, but there are clear limitations. Nonetheless, the graph is largely consistent with our contemporary understanding of mammalian development, despite being constructed through automated procedures. To facilitate its exploration, we created an interactive website in which the nodes and edges shown in **Fig. 1c** can be navigated (https://chengxiangqiu.github.io/tome/).

### Inference of the approximate spatial locations of cell states during mouse gastrulation

Spatial relationships amongst cells are a crucial aspect of development, but this information is lost while profiling disaggregated cells or nuclei. Towards addressing this in part, several groups have developed *in silico* methods for integrating scRNA-seq data with spatially resolved gene expression profiles obtained by fluorescence *in situ* hybridization (FISH) or other means (Satija et al. 2015; Karaiskos et al. 2017). Here we sought to leverage data recently generated by Peng and colleagues, who applied cryosectioning and bulk RNA-seq (GEO-seq) to obtain spatially resolved transcriptomes for precise territories of the mouse embryo from E5.5 to E7.5 (Peng et al. 2019). Inspired by an analysis by Peng *et al*. estimating the regionalization of endodermal subclusters across E7.0 GEO-seq territories, we leveraged TOME to estimate the abundance of individual cell types within each GEO-seq territory (**Fig. 3a**; **Supplementary Fig. 4**; **Supplementary Table 4**) (Newman et al. 2019). For many cell types and territories, this approach appeared to work quite well. For example, the GEO-seq territories inferred to be composed of rostral and caudal neuroectoderm, caudal lateral epiblast, and surface ectoderm are clearly distinguishable at E7.5, in a pattern consistent with expectation (**Fig. 3b**) (Tam and Behringer 1997). Also at E7.5, what we had annotated prior to this analysis as different subsets of paraxial mesoderm (A & B) are also regionalized to the anterior and posterior embryo, respectively (**Fig. 3c**). Finally, we observe the anticipated convergence of embryonic visceral endoderm and definitive endoderm cells during gut development (Nowotschin and Hadjantonakis 2020) (**Fig. 3d**).

**Figure 3.**
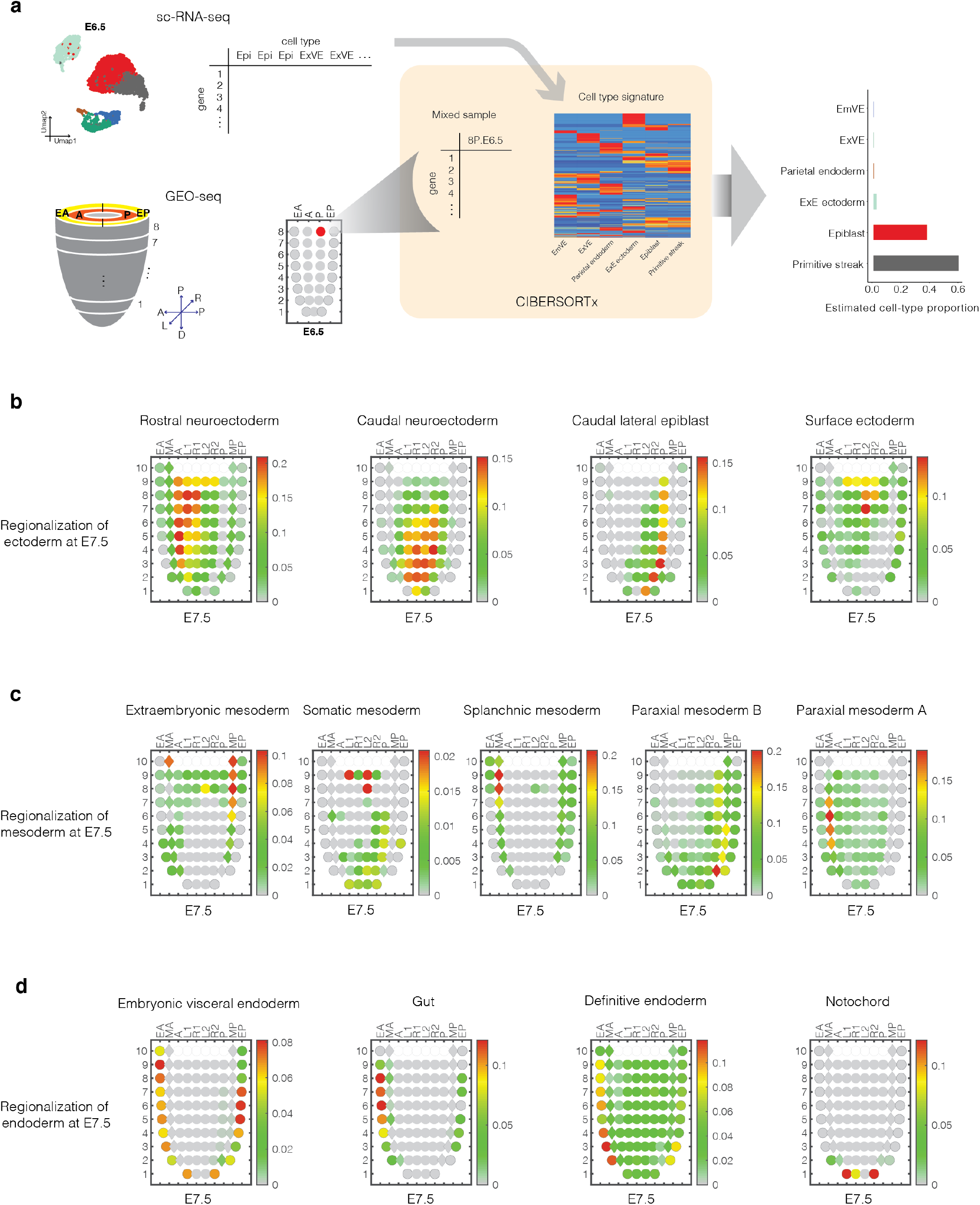
Inference of the approximate spatial locations of cell states during mouse gastrulation. **a,** Inference of cell type contributors to each spatial territory of the gastrulating mouse embryo based on the application of *CIBERSORTx* to GEO-seq data (Newman et al. 2019; Peng et al. 2019). GEO-seq yields bulk RNA-seq data from small numbers of cells dissected from precise anatomic regions of the gastrulating embryo (Peng et al. 2019). For each timepoint for which GEO-seq data was available (E5.5, E6.0, E6.5, E7.0, and E7.5), we estimated a gene expression signature for each cell state from scRNA-seq data, downsampling to 50 cells per state. As E6.0 scRNA-seq data was not available, we instead used data from E6.25 for that timepoint. We then estimated the proportional contribution of each cell state to each GEO-seq sample using *CIBERSORTx* (Newman et al. 2019). ExE: extraembryonic. EmVE: embryonic visceral endoderm. ExVE: extraembryonic visceral endoderm. **b,** Corn plots (Peng et al. 2019) showing the spatial pattern of inferred contributions of various ectodermal cell types at E7.5. **c,** Corn plots showing the spatial pattern of inferred contributions of various mesodermal cell types at E7.5. **d,** Corn plots showing the spatial pattern of inferred contributions of various endodermal cell types at E7.5, as well as notochord. In each corn plot, each circle or diamond refers to a GEO-seq sample, and its weighted color to the estimated cell type composition. Corn plot nomenclature from (Peng et al. 2019). A, anterior; P, posterior; L, left lateral; R, right lateral; L1, anterior left lateral; R1, anterior right lateral; L2, posterior left lateral; R2, posterior right lateral; Epi1 and Epi2, divided epiblast; M, whole mesoderm; MA, anterior mesoderm; MP, posterior mesoderm; En1 and En2, divided endoderm; EA, anterior endoderm; EP, posterior endoderm.

### Inferring the molecular histories of individual cell types

We next sought to infer continuous expression levels for individual genes over the course of each cellular trajectory, focusing on derivatives of the epiblast from E6.25 onwards. First, we leveraged the fact that individual embryos do not correspond precisely to their intended timepoints. Using pseudotime, we ordered the pseudobulk expression profiles of individual embryos (or pools of embryos comprising each sample, in the case of (Pijuan-Sala et al. 2019)). The resulting ordering, which was robust to downsampling, corresponds well with developmental age but may additionally distinguish earlier vs. later individuals/pools at each intended timepoint (**Supplementary Fig. 5a-b**).

Next, for each epiblast-derived cell type that was detectable at E13.5, we calculated a smoothed expression profile along its inferred history, as illustrated in **Supplementary Fig. 5c** for selected genes in one cell type from each germ layer. Despite including the data source as a covariate, these inferred trajectories remained modestly confounded by batch effects across E8.5 → E9.5, *i.e.* the switch from cell-based 10X Genomics data to nucleus-based sci-RNA-seq3 data (**Supplementary Fig. 6**). Nonetheless, at least anecdotally, key transcription factors (TFs) were often sharply upregulated in association with a cell type’s first appearance (**Supplementary Fig. 5c, columns 1 & 2**), while other genes exhibited more gradual patterns of change in relation to pseudotime (**Supplementary Fig. 5c, columns 3 & 4**).

### Systematic nomination of key transcription factors for cell type specification

Inspired by these anecdotal examples, we next sought to leverage TOME to systematically identify TFs that are strong candidates for specifying each newly emerging cell type throughout early mammalian development (Lambert et al. 2018; Niwa 2018). First, we identified 1,391 mouse proteins that are putative TFs, based on orthology with a curated list of human TFs (**Supplementary Table 5**) (“The Human Transcription Factors” n.d.). Then, for each branchpoint in TOME at which a given cell type first emerged, we heuristically defined key TF candidates as those: 1) significantly upregulated in the newly emerged cell type, relative to the pseudo-ancestor; 2) detected in at least 10% of cells in the newly emerged cell type; and 3) not significantly upregulated at any “sister” edges, relative to the newly emerged cell type (**Fig. 4a**). For each such key TF candidate, we calculated a normalized score based on the fold-difference of its expression between the new cell type and its ancestor/sister(s).

**Figure 4.**
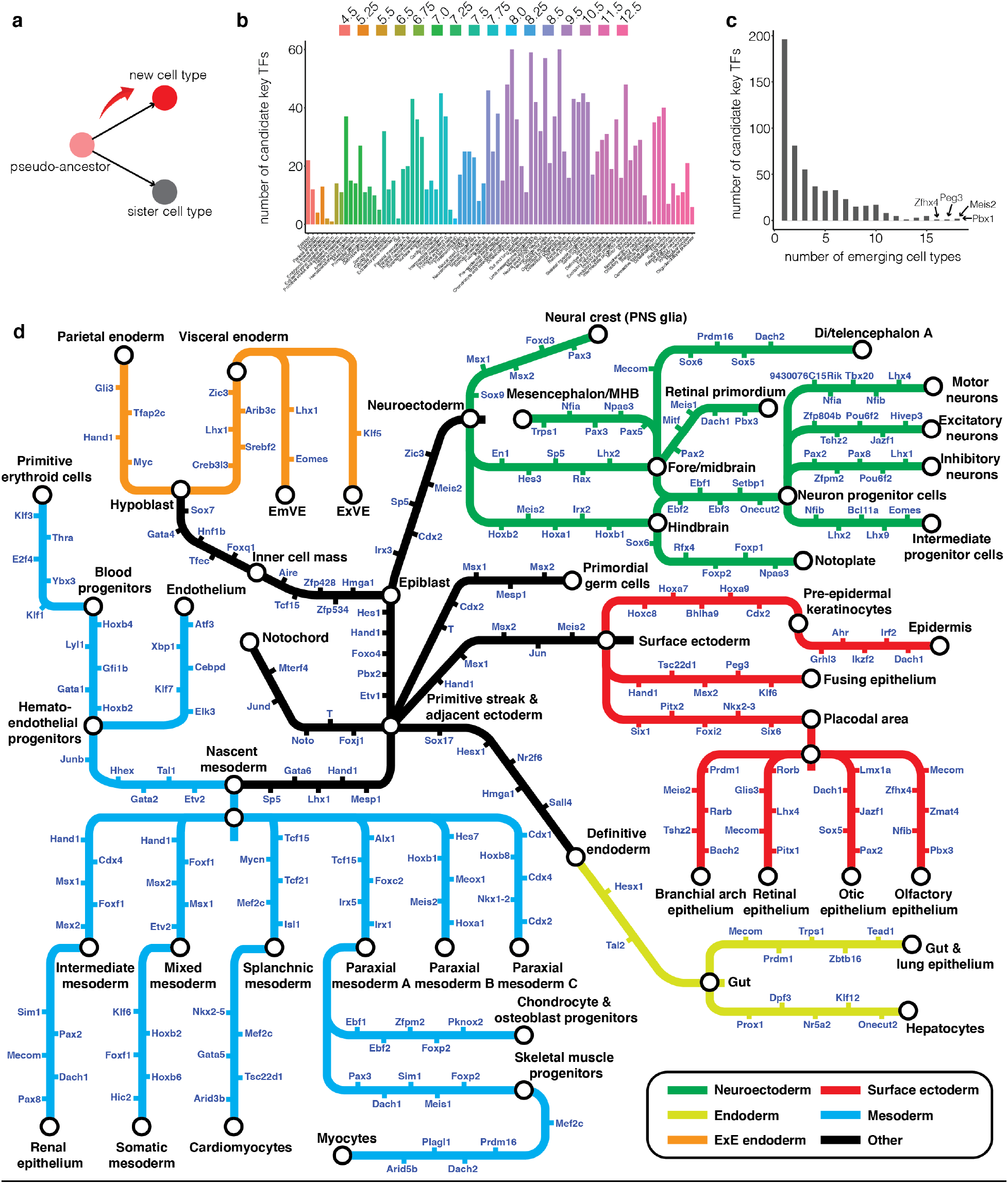
Systematic nomination of candidate key transcription factors for cell type specification. **a,** We heuristically defined candidate key TFs as those that are expressed in the pseudo-ancestral cell state, are significantly upregulated in the newly emerged cell type, are not significantly upregulated at any sister edges. **b,** Histogram of the number of candidate key TFs for each cell type at the timepoint of its first emergence. **c,** The histogram of the number of cell types in which each TF was nominated as a candidate key TF. **d,** Diagram illustrating selected cellular trajectories from TOME, decorated with the top 5 scoring candidate key TFs for each edge. EmVE: embryonic visceral endoderm. ExVE: extraembryonic visceral endoderm. PNS: peripheral nervous system. MHB: midbrain-hindbrain boundary. Style inspired by (Morris et al. 2014).

Altogether, we identified 531 candidate key TFs associated with the emergence of one or more of 82 cell types (24 +/- 15 per cell type; **Fig. 4b**; **Supplementary Table 6**). Most candidate key TFs were specific to one or a few cell types (52% associated with only 1 or 2 cell types). For example, *Gsc* (goosecoid) was identified as a candidate key TF for the emergence of the anterior primitive streak, but no other cell type, and *Srf* for cardiomyocytes, but no other cell type (Blum et al. 1992; Nelson et al. 2005; Miano et al. 2004). On the other hand, a few TFs, such as *Meis2* and *Pbx1*, were associated with the emergence of dozens of cell types (**Fig. 4c**). In **Fig. 4d**, we show the top scoring candidate key TFs for selected trajectories. Despite our automated approach that relied on a handful of datasets, many of these TFs are established as playing critical roles in the emergence of the corresponding cell types. For example, for cardiomyocytes, the top three TFs identified are *Nkx2-5*, *Mef2c*, and *Gata5* (Harvey 1996; Materna et al. 2019; Singh et al. 2010); for notochord, *Foxj1*, *Tbxt* (Brachyury), and *Noto* (Beckers et al. 2007; Herrmann and Kispert 1994; Zizic Mitrecic et al. 2010); for neural crest (PNS glia), *Sox9, Msx1,* and *Msx2* (Cheung and Briscoe 2003; Ishii et al. 2005; Tribulo et al. 2003); and for hematoendothelial progenitors, *Etv2*, *Tal1*, and *Gata2* (Garry 2016; Elcheva et al. 2014; de Pater et al. 2013).

Multilineage priming (MLP) has extensively been documented in hematopoietic lineages and more recently in *C. elegans* (Packer et al. 2019; Laslo et al. 2006). As one form of MLP, we also sought to identify TFs whose *reduced* expression was associated with cell type emergence, which we defined as those: 1) detected in at least 10% of cells in the pseudo-ancestor; 2) significantly downregulated in the newly emerged cell type, relative to the pseudo-ancestor; and 3) both detected in at least 10% of cells and not significantly downregulated at any “sister” edges, relative to the newly emerged cell type. Altogether, we identified 339 candidate key TF whose reduced expression is associated with the emergence of one or more of 76 cell types (13 +/- 12 per cell type; **Supplementary Table 7**). For example, at the split from inner cell mass to epiblast and hypoblast at E4.5, *Gata6* and *Nanog* are identified solely as decreasing in the respective emergence of the epiblast and hypoblast, consistent with the literature (Mitsui et al. 2003; Schrode et al. 2014). Also, *Pou5f1* (Oct4) is identified as a key TF with reduced expression in association with 19 cell types, but increased expression with only 1, consistent with its established role in stemness (**Supplementary Fig. 7a**) (Nichols et al. 1998; Pan et al. 2002). In sharp contrast, *Nfia* and *Nfib* (nuclear factors I/a and I/b) are nominated as key TFs at the emergence of 15 and 14 cell types, respectively, but in all cases upregulated, consistent with broad roles in lineage progression (Chen et al. 2017; Chaudhry, Lyons, and Gronostajski 1997).

### Identification of *cis-*regulatory motifs involved in *in vivo* cell type specification

Although single cell chromatin accessibility profiling (*e.g.* sc-ATAC-seq) is increasingly enabling the ascertainment of *cis*-regulatory programs in embryonic and fetal tissues (Domcke et al. 2020; Cusanovich et al. 2018; Pijuan-Sala et al. 2020), such data is not yet available for a dense timecourse of early development for any of the three species considered here. As a step forward with sc-RNA-seq data alone, we sought to identify DNA sequence motifs that are enriched in the core promoters of developmentally regulated genes in TOME. First, we extended the approach described above to nominate key TFs whose upregulation or downregulation is associated with the emergence of each cell type, to *all* genes. Altogether, this yielded 7,318 key genes associated with the emergence of one or more of 82 cell types (345 +/- 246 per cell type; **Supplementary Fig. 7b**; **Supplementary Table 8**). Second, for each cell type, we applied *HOMER* (Heinz et al. 2010) to discover DNA sequence motifs that are specifically enriched in the core promoters of key genes (-300 to +50 bp of annotated TSSs). Finally, we estimated q-values for discovered motifs by data label permutation. At an FDR of 10%, we implicated 77 *de novo* promoter motifs in the emergence of 41 mouse cell types (**Supplementary Table 9**), as well as an additional 100 previously documented promoter motifs (some overlapping with the *de novo* set) in the emergence of 30 mouse cell types (**Supplementary Table 10**).

We then asked whether the sequence motifs identified in the core promoters of developmentally regulated genes correspond to the binding sites of candidate key TFs for the same cell types, which would provide a plausible confirmation of their role. We identify 20 such instances, 15 of which are positive correlations (*i.e.* consistent directionality between TF expression and target gene expression) and 5 of which are negative correlations (**Supplementary Table 11**). For example, the transcriptional activator Rfx3 is sharply upregulated at the emergence of the notochord at E7.25, and its cognate motif is strongly enriched at the promoters of key genes upregulated in these same cells (**Supplementary Fig. 8a-c**) (Beckers et al. 2007; Bonnafe et al. 2004). In contrast, the transcriptional repressor Snail, encoded by *Snai1*, is upregulated at the emergence of nascent mesoderm at E6.75, but its cognate motif strongly enriched in the promoters of downregulated key genes (**Supplementary Fig. 8d-f**) (Hemavathy, Ashraf, and Ip 2000; Carver et al. 2001). Of note, RFX3 motifs are very strongly enriched near the TSSs of notochord-upregulated genes, while SNAIL1 motifs are more diffusely enriched across the promoters of nascent mesoderm-downregulated genes (**Supplementary Fig. 8b, e**).

A limitation of these analyses is that we restricted our search for enriched sequence motifs to the core promoters of up- or down-regulated key genes. As single cell, genome-wide chromatin accessibility datasets spanning mammalian embryogenesis become available, such analyses can be extended to enhancer-mediated regulation.

### Systematic comparison of the cellular trajectories of mouse, zebrafish and frog embryogenesis

The origins and evolution of vertebrate cell types are fascinating topics on which the single cell profiling of embryogenesis may shed much needed light (Arendt et al. 2016). However, even if we adopt an evolutionary definition of cell types, it remains unclear how best to identify “cell type homologs” across vast evolutionary distances. To facilitate the systematic alignment of cell types across vertebrates, we applied the same strategy used for TOME to zebrafish (*D. rerio*) and frog (*X. tropicalis*) embryogenesis, again relying on publicly available single cell RNA-seq datasets. For zebrafish, we integrated data from two studies that used different technologies but together included 15 developmental stages, beginning at the high stage (hpf 3.3) and ending at the early pharyngula stage (hpf 24), essentially spanning epiboly and segmentation (**Fig. 5a**; **Supplementary Table 1**) (Wagner et al. 2018; Farrell et al. 2018). The resulting graph contains 221 nodes, each assigned one of 63 cell type annotations, and 257 edges with weights greater than 0.2 (**Fig. 5b**). Marker genes used to annotate cell types are provided in **Supplementary Table 12**, and all edge weights in **Supplementary Table 13**. We also nominated key upregulated and downregulated TFs using the same approach described for mouse development above (**Supplementary Tables 14-15**).

**Figure 5.**
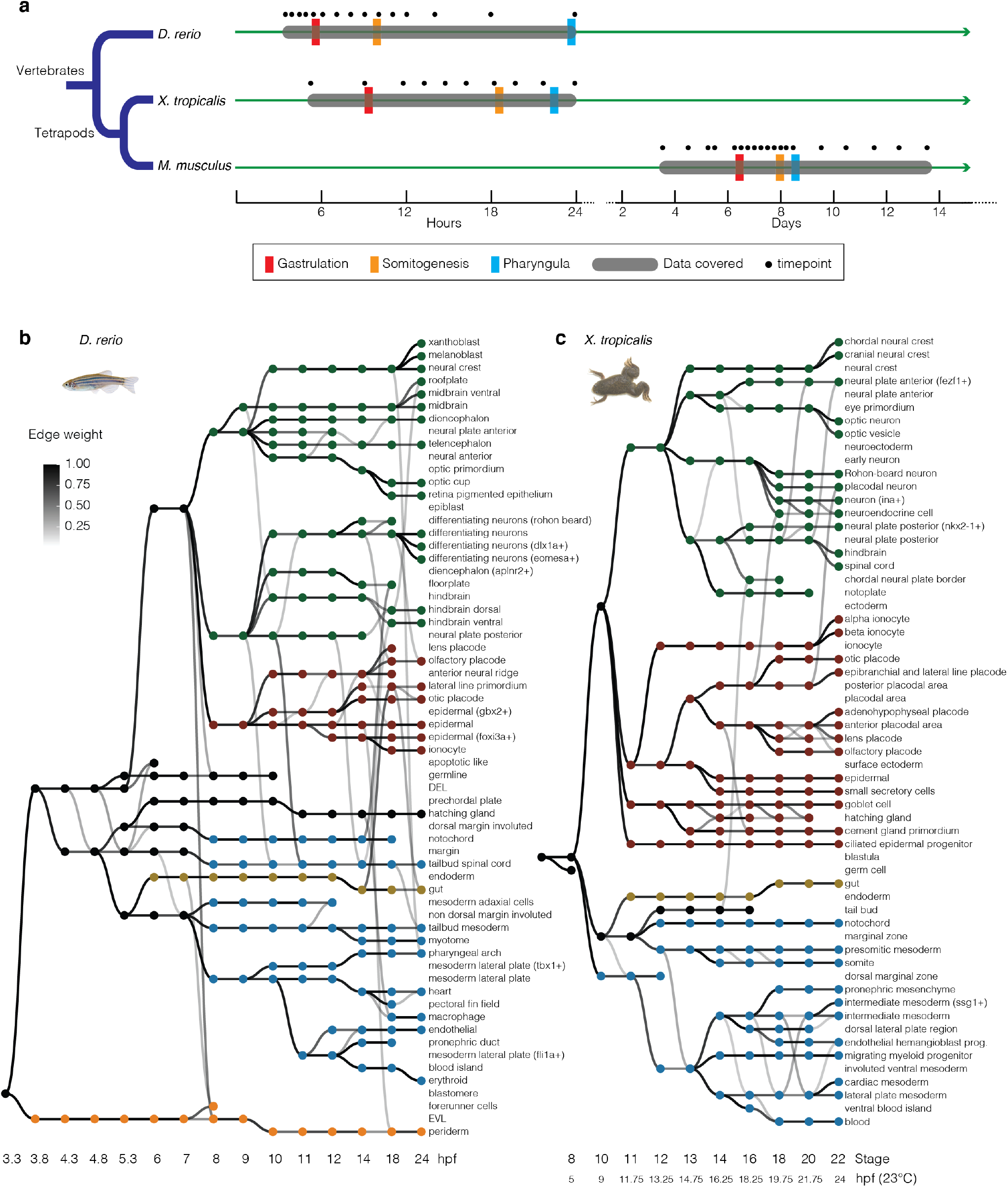
Reconstruction of the cellular trajectories of zebrafish and frog embryogenesis. **a,** Comparative developmental timelines for mouse, zebrafish, and frog, spread over two time scales, and approximate (as may depend on temperature, particularly for frog). “Gastrulation” and “Somitogenesis” refer to the timing of onset of these processes (Afonin et al. 2006). “Pharyngula” refers to the timing of onset of pharyngeal arch formation (Irie and Kuratani 2011). Black dots refer to timepoints sampled across seven studies. Grey rounded rectangles indicate developmental windows covered by cellular trajectory reconstructions. **b,** Directed acyclic graph showing inferred relationships between cell states across early zebrafish development. Each row corresponds to one of 63 cell type annotations, and columns to developmental stages spanning hpf3.3 to hpf24. Nodes denote cell states, and node colors denote germ layers. All edge weights greater than 0.2 are shown in grey-scale. **c,** Directed acyclic graph showing inferred relationships between cell states across early frog development. Each row corresponds to one of 60 cell type annotations, columns to developmental stages spanning S8 (hpf5, 23C) to S22 (hpf24, 23C), nodes to cell states, and node colors to germ layers. All edge weights greater than 0.2 are shown in grey-scale. DEL: deep cell layer. EVL: enveloping layer.

For frog, we re-analyzed one dataset spanning 10 developmental stages, from S8 and S22 (Briggs et al. 2018), spanning gastrulation and neurulation (**Fig. 5a**; **Supplementary Table 1**). The resulting graph contains 192 nodes, each assigned one of 60 cell type annotations, and 221 edges with weights greater than 0.2 (**Fig. 5c**). Marker genes used to annotate cell types are provided in **Supplementary Table 16**, all edge weights in **Supplementary Table 17**, and candidate key TFs in **Supplementary Tables 18-19**.

We next sought to systematically align cell types from each species to their “cell type homologs” in the other two species. Because *M. musculus* is separated from *D. rerio* and *X. tropicalis* by ∼450 million and ∼360 million years of evolution, respectively, these alignments proved much more challenging than integrating data from more closely related species such as mouse and human (Cao et al. 2020; Yu et al. 2019). We attempted three strategies.

As a first strategy, treating cells of each state from each timepoint as a “pseudo-cell”, we integrated data from all three species with anchor-based batch correction (Stuart et al. 2019). Within the resulting UMAP co-embedding of 765 pseudo-cells, we could identify 15 major groups — epiblast, early gastrulation, neuroectoderm, surface ectoderm, mesoderm, floor plate & roof plate, gut, placodal area, neural crest, epithelium, brain, neurons, endothelium, myocytes, and blood — each containing cell states from all three species (**Supplementary Fig. 9**). However, within each such major group, the homology between specific cell types generally remained ambiguous.

As a second strategy, we performed all possible pairwise comparisons between the transcriptomes of cell types of each pair of species, excluding extraembryonic lineages (Fukazawa et al. 2010). First, we performed cell type correlation analysis (Cao et al. 2019), which uses a regression framework to ask, between each pair of species, which cell types are the best reciprocal best matches to one another (**Supplementary Fig. 10**; **Supplementary Table 20**; **Methods**). We then manually reviewed the highest ranking cell type pairings for biological plausibility. For mouse vs. zebrafish, out of 4,543 pairings tested, 147 were highly ranked, of which we selected 44 as the most biologically plausible (**Supplementary Table 21**). Exclusion criteria included the cell types arising from different germ layers or major groups (as defined in **Supplementary Fig. 9**), arising at very different temporal stages, or if a cell type was exclusive to one species. In cases where multiple related matches were observed, we generally selected the match with the highest *β* score. Applying this same approach to mouse vs. frog and zebrafish vs. frog, we identified 20 and 47 plausible cell type homologs pairings, respectively (**Supplementary Fig. 11a**; **Supplementary Table 21**).

As a third strategy, we focused on overlaps between the candidate key TFs associated with the emergence of each cell type in each species. For each possible interspecies pairing of cell types, we identified orthologous TFs that were nominated in both, and then adopted a permutation approach to identify instances in which an excess of orthologous candidate key TFs were shared between the cell types. For mouse vs. zebrafish, out of 4,312 pairings tested, 88 exhibited more sharing than >99% of permutations, of which we retained 26 as the most biologically plausible (**Supplementary Table 22**). Applying this same approach to mouse vs. frog and zebrafish vs. frog, we identified 19 and 19 plausible cell type homolog pairings, respectively (**Supplementary Fig. 11b**; **Supplementary Table 22**).

Some candidate cell type homologs overlapped between these second and third strategies (**Fig. 6a**; **Supplementary Table 23**). Overall, we were able to assign at least one cell type homolog to 48 of 77 embryonic mouse cell states, 52 of 59 zebrafish embryonic cell states, and 44 of 60 frog embryonic cell states. Some loosely annotated cell types were resolved through homology. For example, zebrafish *eomesa*+ and *dlx1a*+ differentiating neurons were homologous to mouse intermediate progenitor cells and inhibitory interneurons, respectively. In certain cases, we observed “three way” pairwise homology and nominated regulators (**Fig. 6b**). For example, *Gsc*, a canonical TF of the Spemann organizer (De Roberts et al. 1992), was nominated as a key regulator of the anterior primitive streak (mouse), dorsal margin involuted (zebrafish), and dorsal marginal zone (frog), cell types that were also identified as homologs of one another. Other such “three way” nominated TF regulators and associated cell types include *Sox7* for haemogenic endothelium (Costa et al. 2012)*, Tbx2* for the otic placode (Takabatake, Takabatake, and Takeshima 2000; Barrionuevo et al. 2008) and *Gata3* for surface ectoderm (**Fig. 6b**).

**Figure 6.**
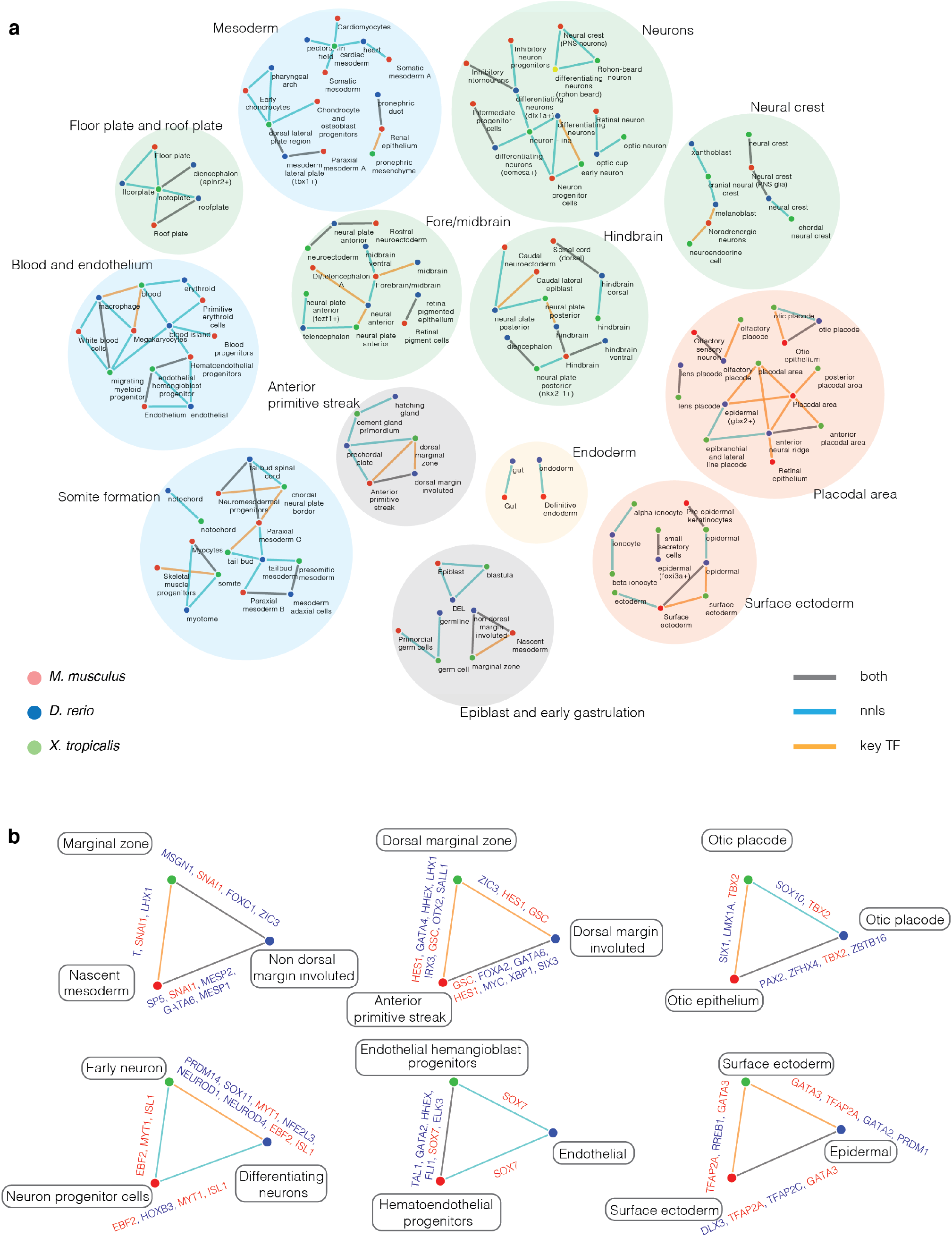
The union of candidate cell type homologs identified between three species (mouse, zebrafish, frog) by two strategies. **a,** Candidate cell type homologs were identified either by comparison of transcriptomes via non-negative least squares (“nnls”) regression or by examining overlap between upregulated candidate key TFs (“key TF”). Nominated pairings were manually reviewed, and a subset retained based on biological plausibility. Colors of nodes indicate the species of a given cell type, and colors of edges indicate which approach(es) identified the pairing. Sets of connected candidate cell type homologs are further grouped by germ layer or developmental system. PNS: peripheral nervous system. DEL: deep cell layer. **b,** Selected examples of “three way” pairwise cell type homology from different germ layers in the above network. Upregulated candidate key TFs shared by each pair of species are listed, with the subset shared by all three species in red font.

## Discussion

Nearly forty years ago, Sulston and colleagues painstakingly mapped out the entirety of the invariant embryonic cell lineage of *C. elegans*, comprising 671 cells (Sulston et al. 1983). The Sulston map provided a foundational scaffold for the integration of future experimental results, as well as a precise nomenclature for the discussion of specific subsets of cells within the developing worm. Recently, Packer and colleagues intersected the Sulston lineage with the mRNA profiles of the same cells, shedding fresh light on the relationship between cell states and fates (Packer et al. 2019).

Can equivalently global views of development be achieved for the developing mouse? For reasons including scale, complexity, variance and accessibility, this is an extraordinary challenge and one that may take decades to fully come to fruition, if indeed it ever does. However, given the pace at which relevant technologies are emerging and evolving, it feels increasingly tractable to make meaningful progress.

Here, towards a scaffold for such an undertaking, we sought to leverage recently published single cell RNA-seq data to construct a “roadmap” of molecular trajectories that cells traverse during the peri-implantation, gastrulation and organogenesis stages of mouse development (**Fig. 1**). Our approach — constructing a directed acyclic graph wherein each node corresponds to a group of related cells at a given timepoint, and each edge to similarity between groups observed at adjacent timepoints — is highly reductionist. However, we believe that this framing provides a useful entry point for analyses that benefit from a global view of developmental processes. For example, in addition to systematically nominating specific TFs as key regulators of the initial emergence of each cell type, we are able to assess which TFs and genes appear to have relatively specific vs. general roles in development (**Fig. 4**; **Supplementary Fig. 7**). Furthermore, by constructing developmental graphs for additional vertebrate species through the same method, we can identify “cell type homologs” through approaches that consider all cell types in each pair of species, analogous to the comparison of genomes (**Figs. 5, 6**).

Although “cell type” is a useful concept, a limitation of this terminology is that it obscures continuous aspects of heterogeneity — *e.g.* as might be expected for spatial gradients or during the maturation of a cell type. This framing also forces us to make decisions about the level of resolution at which to define cell types. Although we have made some progress in relating TOME to spatial information (**Fig. 3**), new nomenclature that facilitates the discussion of precise subsets of cells within spatially or otherwise heterogeneous cell types is sorely needed.

As discussed above, molecular trajectories are not equivalent to cellular phylogenies, but are likely to constrain them. Indeed, the cell type relationships inferred here on the basis of gene expression are largely consistent with our understanding of the *bona fide* ancestors and descendants of cell types in mouse development. Exceptions and ambiguities may represent errors that can be clarified through deeper analysis (**Fig. 2**), or potentially novel relationships that require validation. To that end, *in vivo,* organism-scale lineage recording, originally developed in zebrafish, has recently been adapted to the mouse (McKenna et al. 2016; Bowling et al. 2020; Kalhor et al. 2018; Chan et al. 2019). Although such methods remain very far from delivering anything approaching the resolution of the Sulston lineage, they can likely be applied in their current form to support or reject potential lineage relationships suggested here. Additionally, even within cell states, such lineage data might shed light on the patterns of cell division that underlie the differentiation and proliferation of each cell type.

TOME provides a scaffold onto which additional single cell gene expression datasets from mouse development can be layered. In this vein, the remarkable consistency and robustness of *in vivo* mouse development is a terrific feature, and contrasts with *in vitro* differentiation time courses, which may vary by lab, operator, cell line, etc. In terms of additional data generation, near-term goals that we are pursuing include consistent sampling throughout mouse development at 6 hour intervals, better integration across the E8.5-E9.5 “technology switch”, and extending whole embryo scRNA-seq datasets out to P0. We also anticipate additional single cell data types (*e.g.* chromatin accessibility, methylation, histone modifications, transcription factor binding, etc.) can be generated by independent groups and/or technologies and layered onto TOME as well. We are particularly excited about the possibility of linking the temporal unfolding of combinatorial TF expression to enhancer accessibility, and then enhancer accessibility to the expression of *cis*-regulated genes.

## Methods

### Systematic reconstruction of the cellular trajectories of mouse embryogenesis

Single cell or single nucleus RNA-seq data were collected from four studies spanning 19 timepoints between E3.5 and E13.5 of mouse embryogenesis, collectively 1,443,229 cells from 468 samples, where each sample consists of either a single mouse embryo or a small pool of embryos from the same timepoint (Mohammed et al. 2017; Cheng et al. 2019; Pijuan-Sala et al. 2019; Cao et al. 2019). The details are summarized in **Supplementary Table 1**. For each dataset, we took the unique molecular identifiers (UMI) count matrix (feature ✕ cell) from the data source and separated cells by timepoint. For each timepoint, we performed conventional single-cell RNA-seq data processing using *Seurat*/v3: 1) normalizing the UMI counts by the total count per cell followed by log-transformation; 2) selecting the 2,500 most highly variable genes and scaling the expression of each to zero mean and unit variance; 3) applying PCA and then using the top 30 PCs to create a *k*-NN graph, followed by Louvain clustering (resolution = 1); 4) performing UMAP visualization in 2D space (dims = 1:30, min. dist = 0.75) (Stuart et al. 2019). For some timepoints, we observed obvious batch effects with respect to either study or sample identity. We therefore performed an additional batch correction before the PCA, following the standard pipeline for dataset integration in *Seurat*/v3 (https://satijalab.org/seurat/v3.2/integration.html), using either the study or sample identity to split datasets, followed by identifying “anchors” between pairs of post-splitting subsets of the datasets (features = 2500, k.filter = 200, dims = 1:30) (**Supplementary Fig. 1c-d**).

For cell clustering, we manually adjusted the resolution parameter towards modest overclustering, and then manually merged adjacent clusters if they had a limited number of differentially expressed genes (DEGs) relative to one another (for this purpose, DEGs were defined as genes expressed at mean >0.5 UMIs per cell across the pair of clusters with a >4-fold difference between the clusters) or if they both highly expressed the same literature-nominated marker genes. Subsequently, we annotated individual cell clusters using 2-5 literature-nominated marker genes per cell type label (**Supplementary Table 2**). Most of the cell type labels and associated marker genes were obtained from the four studies that generated the data. However, we double-checked each cell type assignment, often with additional marker genes. Importantly, we revisited and revised some of the cell type or trajectory annotations of (Cao et al. 2019), *e.g.* “Ependymal cell” → “Roof plate”; “Isthmic organizer cells” → “Mesencephalon/MHB”. A full list of these annotation revisions is provided in **Supplementary Table 24**.

To connect each cell state observed at a given timepoint with its “pseudo-ancestors”, we first merged all cells from that timepoint and the preceding timepoint using *Seurat*/v3. Integration and batch correction were performed as described above, except that we also split based on timepoint identity (features = 2500, k.filter = 200, dims = 1:30). Because of the very large number of cells, we used a reciprocal PCA-based space (Stuart et al. 2019) to find anchors for pairs of timepoints that included data from (Cao et al. 2019). After integration, we performed PCA and then used the top 30 PCs to co-embed cells as a 3D UMAP (min. dist = 0.75), from which we calculated Euclidean distances between individual cells from the earlier and later timepoints.

We then determined edge weights between cell states of the successive timepoints using a bootstrapping strategy. For cells of each cell state at the later timepoint, we identified their five closest neighbor cells from the earlier timepoint and then calculated the proportion of these neighbors derived from each potential antecedent cell state. We repeated these steps 500 times with 80% subsampling from the same embedding. We then took the median proportions as the set of weights for edges between a cell state and its potential antecedents. To evaluate the robustness of this approach to the choice of co-embedding space, we repeated it using Euclidean distances between cells in PCA space (dims = 30) instead of UMAP space (dims = 3). The resulting edge weights were highly correlated (Pearson correlation coefficient = 0.987). We evaluated the above approach with *k* parameters (for the *k-*NN) other than five, and found the resulting edge weights to be highly correlated with those obtained with *k* = 5 (Pearson correlation coefficients from 0.998 to 0.999 for *k* = 8, 10, 15, 20). Edge weights > 0.2 from the UMAP embedding were retained for the resulting acyclic directed graph shown in **Fig. 1c**.

We repeated this strategy to generate similar graphs for zebrafish (*D. rerio*) and frog (*X. tropicalis*) embryogenesis, again relying on publicly available scRNA-seq datasets. For zebrafish, we integrated data from two studies that overlapped at three timepoints (hpf6, hpf8, hpf10); we excluded cells from hpf4 because of excessive batch effects (Wagner et al. 2018; Farrell et al. 2018). For frog, we used cells from a single study (Briggs et al. 2018). Further details regarding data sources are available in **Supplementary Table 1**.

### RNA velocity analysis

Two datasets were used for performing RNA velocity analysis (Mohammed et al. 2017; Cheng et al. 2019; Pijuan-Sala et al. 2019; Cao et al. 2019). For the Pijuan-Sala *et al*. dataset, which was generated on the 10X Genomics platform, we downloaded the raw data (E-MTAB-6967) and processed it using *kb-python* (Melsted et al. 2019). For the Cao *et al*. dataset, which was generated with sci-RNA-seq3, we processed the raw data using the basic sci-RNA-seq pipeline followed by extracting the spliced reads and unspliced reads for each cell using *velocyto* (La Manno et al. 2018; Cao et al. 2019). The RNA velocity analysis and UMAP visualization were performed with *Scanpy*/v.1.6.0 and *scVelo* (Bergen et al. 2020; Wolf, Angerer, and Theis 2018). Briefly, genes with low expression were filtered out (min_counts = 5, min_counts_u = 5), and each cell’s counts were normalized towards the median UMI counts per cell by a scaling factor. The 3000 genes with the highest variance were selected, and the data was log-transformed after adding a pseudocount. The spliced and unspliced count matrices were similarly filtered and normalized. We then applied *scvelo.pp.memoments* and *scvelo.tl.velocity* for velocity estimation (n_pcs = 30, n_neighbors = 30), followed by *scvelo.tl.velocity_graph* and *scvelo.tl.umap* for data visualization (min_dist = 0.75).

### Inferring the molecular histories of individual cell types

For this particular analysis, because one dataset did not include the extraembryonic tissues (Cao et al. 2019), we excluded cells annotated as derived from the extraembryonic lineages (embryonic visceral endoderm, extraembryonic visceral endoderm, parietal endoderm, and extraembryonic ectoderm). For E6.5, the sequencing depths were very different between datasets, so we only used cells from the Pijuan-Sala *et al*. dataset. In addition, the Pijuan-Sala *et al*. dataset pooled multiple embryos per sample, so we used sample identity instead of embryo identity. In the end, four samples from the Cheng *et al*. dataset, 34 samples from the Pijuan-Sala *et al*. dataset, and 61 samples from the Cao *et al*. dataset were used for the pseudobulk analysis. UMI counts mapping to each sample were aggregated to generate a pseudobulk RNA-seq profile for each sample. We then applied the *fit_models* function of *Monocle/3* to identify genes that were highly correlated with the embryos’/samples’ staged age (model_formula_str = ∼stage + dataset). To mitigate major batch effects between cell vs. nucleus-derived subsets of the data, we separately performed DEG analysis on the samples from before and including E8.5 (n = 38) vs. after E8.5 (n = 61), and then took the union of the top 3,000 genes with the lowest *q* values identified in each subset. We then filtered out genes that were significantly different between the pre- and post-E8.5 subsets (p_value < 0.05). This left 865 genes, which were used to construct a pseudotime trajectory using *DDRTree* as implemented in *Monocle/*v*2* (Qiu et al. 2017). Each embryo/sample was assigned a pseudotime value on the basis of its position along the trajectory. Of note, this ordering was highly robust to 80% subsampling (all Pearson correlation coefficients were >0.99 between pseudotimes derived from 100 iterations of 80% subsampling vs. the full dataset).

### Deconvolution of cell composition of GEO-seq sample using *CIBERSORTx*

This analysis was performed by running deconvolution on each GEO-seq sample using *CIBERSORTx* with default parameters (Newman et al. 2019; Peng et al. 2019). GEO-seq samples were collected from distinct spatial positions in the mouse embryo with mixed cell populations from E5.5, E6, E6.5, E7, and E7.5 (Peng et al. 2019). For each stage, we first learned a gene expression signature for each cell state at the corresponding timepoint by downsampling it to 50 cells and then summing these. We excluded ExE ectoderm from this set because the GEO-seq experiments only focused on the cell lineages derived from the inner cell mass. Also, because single cell profiles from E6 were missing from the scRNA-seq data integrated here, we used data from E6.25 instead.

### Systematic nomination of key transcription factors for cell type specification

The list of 1,391 mouse proteins that are putatively TFs was based on liftover from a curated list of human transcription factors (http://humantfs.ccbr.utoronto.ca/) (**Supplementary Table 5**). The liftover was done with *BioMart* (Ensembl Genes 102) (Yates et al. 2020). For each edge in TOME at which a given cell type first emerged, we used three criteria to identify key TF candidates: 1) its expression significantly increased in the newly emerged cell type, relative to the pseudo-ancestral cell state (*Seurat*/v3; p_val_adj < 0.05, non-parametric Wilcoxon rank-sum test); 2) it was significantly more highly expressed in the newly emerged cell type, relative to its “sister” edges deriving from the same pseudo-ancestor (by the same test and threshold); 3) it was detected in at least 10% of cells of the newly emerged cell type. For each such candidate key TF, we scaled its log fold-change calculated by either criteria #1 or #2 to unit variance and zero mean (across the set of candidate key TF identified for a given newly emerged cell type) and then averaged these scaled fold-change values to determine a score intended to convey its importance relative to other candidate key TFs for the same cell type.

To identify TFs whose reduced expression was associated with the emergence of each cell type, we looked for those that: 1) are detected in at least 10% of cells of the pseudo-ancestral cell type; 2) are significantly downregulated in the newly emerged cell type, relative to the pseudo-ancestor (*Seurat*/v3; p_val_adj < 0.05, non-parametric Wilcoxon rank-sum test); and 3) are both detected in at least 10% of cells and significantly more highly expressed at “sister” edges, relative to the newly emerged cell type (by the same test and threshold).

The list of 1,183 zebrafish TFs and 1,014 frog TFs was based on liftover from a curated list of human transcription factors (**Supplementary Table 5)**. The liftover for zebrafish TFs was done with *BioMart* (Ensembl Genes 102) (Yates et al. 2020). The liftover for frog TFs was done as part of the original study (Briggs et al. 2018). Candidate key TFs for each cell type emergence in these species were identified and scored as described above for mouse.

### Co-embedding of cell states from three species

We first created a list of homologous genes across the three species by liftover of all gene identities from the three species to the corresponding human gene identities, either based on *BioMart* (Ensembl Genes 102) (Yates et al. 2020) or the original study in the case of frog (Briggs et al. 2018). A list of 23,489 genes was compiled, wherein each of the genes was homologous in at least two species. To create the transcriptional features of each cell state, we first averaged cell-state-specific UMI counts, normalized by the total count, multiplied by 100,000 and natural-log-transformed after adding a pseudocount. We then divided all the cell states from three species into four groups: the mouse single-cell group (n = 142), the mouse single-nucleus group (n = 226), the zebrafish group (n = 205), and the frog group (n = 192). We treated each cell state as a pseudo-cell, performing the anchor-based batch correction approach implemented by *Seurat*/v3 (nfeatures = 5,000, k.filter = 100, dims = 1:30, min.dist = 0.6) (Stuart et al. 2019). For cell states spanning multiple timepoints, cells from each timepoint were treated as a separate pseudo-cell for the purposes of this analysis.

### Identification of interspecies correlated cell types using non-negative least-squared (*NNLS*) regression

We first created a list of homologous genes between each pair of species (n = 13,448 for *mm* vs. *zf*, n = 14,252 for *mm* vs. *xp*, and n = 13,326 for *zf* vs. *xp*), either based on *BioMart* (Ensembl Genes 102) (Yates et al. 2020) or the original study in the case of frog (Briggs et al. 2018). To identify correlated cell types between each pair of species, we first calculated an expression value for each gene in each cell type by averaging the log-transformed normalized UMI counts of all cells of that type across all timepoints at which the cell type appeared. Extraembryonic cell types (inner cell mass, hypoblast, parietal endoderm, extraembryonic ectoderm, visceral endoderm, embryonic visceral endoderm, and extraembryonic visceral endoderm for the mouse; blastomere, EVL, periderm, forerunner cells for the zebrafish) were excluded from this analysis. For mouse E6.5, we only used cells from a single study (Pijuan-Sala et al. 2019). For each pair of species, we took homologous genes and applied non-negative least squares (*NNLS*) regression to predict gene expression in target cell type (*Ta*) in dataset A based on the gene expression of all cell types (*Mb*) in dataset B: *Ta = β0a + β1aMb*, based on the union of the 1,200 most highly expressed genes and 1,200 most highly specific genes in the target cell type. We then switched the roles of datasets A and B, *i.e.* predicting the gene expression of target cell type (*Tb*) in dataset B from the gene expression of all cell types (*Ma*) in dataset A: *Tb = β0b + β1bMa*. Finally, for each cell type *a* in dataset A and each cell type *b* in dataset B, we combined the two correlation coefficients: *β* = 2(*βab* + 0.001)(*βba* + 0.001) to obtain a statistic for which high values reflect reciprocal, specific predictivity.

To identify candidate cell type homologs, we manually reviewed pairings with a *β* score > 1e-4 and that ranked highly from the perspective of both species, *i.e.* where cell type B was one of the top five matches for cell type A and vice versa. We next performed a manual selection based on the following criteria: 1) excluding pairs of cell types which derive from different germ layers or major groups (**Supplementary Fig. 9**) (*e.g.* blood progenitors (*mm*) vs. optic cup (*zf*)); 2) excluding pairs of cell types which emerged at very different temporal stages (*e.g.* rostral neuroectoderm (*mm*) vs. DEL (*zf*)); 3) excluding cell types only expected in one species or the other (*e.g.* hatching gland (*zf*) is not expected in mouse); 4) for cell types which were correlated with multiple cell types with ancestor-descendant relationships in the other species, we selected the one which was more ancestral (*e.g.* hindbrain (*mm*) was correlated with both hindbrain ventral (*zf*) and hindbrain (*zf*), and we assigned it to hindbrain (*zf*)); 5) for cell types which were correlated with multiple cell types in the other species that lacked a clear ancestor-descendant relationship, we selected the pair with the highest *β* score. The details of manual selection are provided in **Supplementary Table 21**.

### Identification of correlated cell types between species based on overlapping key TF candidates

For each possible interspecies pairing of cell types, we identified orthologous TFs that were nominated in both and then calculated, as an estimate of relative likelihood, the product of the frequencies in which each of these TFs were nominated as key in their respective species (to account for the fact that some TFs are nominated in many cell types and therefore more likely to overlap; **Fig. 4c**). To identify which such instances were potentially significant, we repeated these procedures after taking random samples of key TFs without replacement (10,000 times) and retained pairings with estimated relative likelihoods more extreme than 99% of permutations. We then performed a similar manual selection, details of which are provided in **Supplementary Table 22**.

### Identification of *cis-*regulatory motifs involved in *in vivo* cell type specification

As a first step towards identifying *cis-*regulatory motifs involved in cell type identification, we extended to all genes the approach described above to nominate key TFs whose upregulation or downregulation is associated with the emergence of each cell type. For each edge in TOME at which a given cell type first emerged, we used three criteria to identify key gene candidates: 1) its expression significantly increased in the newly emerged cell type, relative to the pseudo-ancestral cell state (*Seurat*/v3; p_val_adj < 0.05, non-parametric Wilcoxon rank-sum test); 2) it was significantly more highly expressed in the newly emerged cell type, relative to its “sister” edges deriving from the same pseudo-ancestor (by the same test and threshold); 3) it was detected in at least 10% of cells of the newly emerged cell type. To identify genes whose reduced expression was associated with the emergence of each cell type, we looked for those that: 1) are detected in at least 10% of cells of the pseudo-ancestral cell type; 2) are significantly downregulated in the newly emerged cell type, relative to the pseudo-ancestor (*Seurat*/v3; p_val_adj < 0.05, non-parametric Wilcoxon rank-sum test); and 3) are both detected in at least 10% of cells and significantly more highly expressed at “sister” edges, relative to the newly emerged cell type (by the same test and threshold).

We used *HOMER*/v4.11 (Heinz et al. 2010) to discover DNA sequence motifs that are specifically enriched in the core promoters of key genes (-300 to +50 bp of annotated TSSs). Running the *findMotifs.pl* function with default parameters, each test set was defined as the core promoters of either upregulated or downregulated key genes at specific cell edges (excluding sets with fewer than 5 key genes), and the background as core promoters of key genes from all edges not in the test set. Motif quality was evaluated based on a q-value, which was calculated for each motif by 100 iterations of randomizing data labels and re-running *HOMER*. Motifs were aligned to known motif binding sequences based on the *JASPAR* and internal *HOMER* databases with default parameters as well (Khan et al. 2018). Mapping of specific motif positions around the TSS was assessed with the *HOMER* function *annotatePeaks.pl* using the following parameters: tss mm10 -hist 10 -ghist.

## Acknowledgments

We thank the members of the Shendure and Trapnell labs for helpful discussions. This work was funded by Paul G. Allen Frontiers Foundation (Allen Discovery Center grant to J.S. and C.T.), the National Institutes of Health (UM1HG011531 to W.S.N., J.S., and C.M.D., R01 NS109425 to C.B.M.), the Bonita and David Brewer Fellowship (to C.Q.). J.S. is an investigator of the Howard Hughes Medical Institute.

## Author Contributions

J.S. and C.Q. designed the research. C.Q. and T.L. performed computational analysis. J.C., X.H., D.C., S.S., W.S.N., and C.T. assisted with data analysis. M.S., C.M.D., and C.B.M. assisted with results interpretation. J.S. and C.Q. wrote the paper, with input from all authors.

---

**Supplementary Figure 1.**
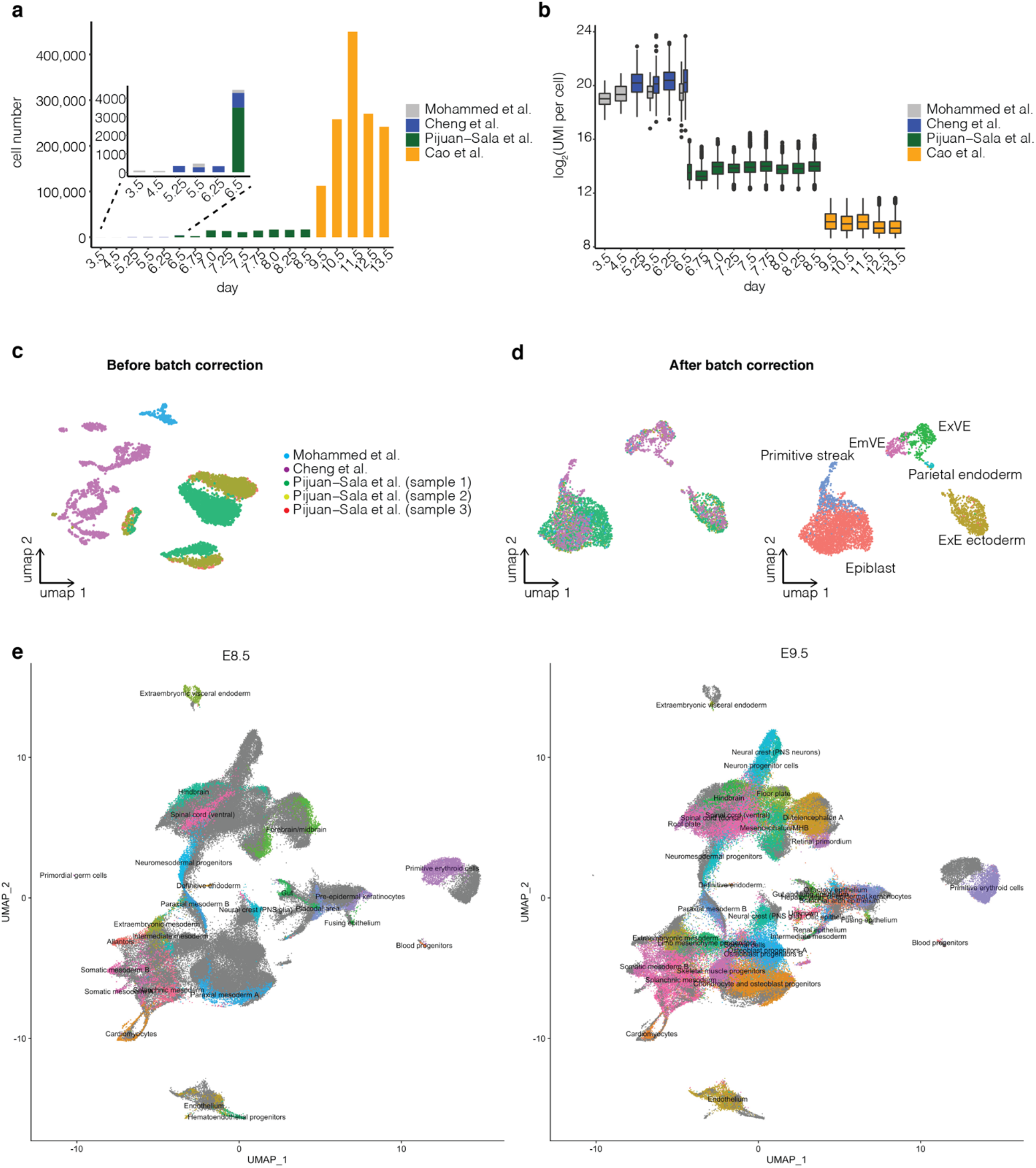
Integration of datasets spanning E3.5 to E13.5 of mouse development. **a,** The number of cells per stage obtained from four studies that collectively span E3.5 to E13.5 of mouse development (Mohammed et al. 2017; Cheng et al. 2019; Pijuan-Sala et al. 2019; Cao et al. 2019). **b,** Box plot of log2(UMI counts) per cell across the stages and studies. **c,** As illustrated by a UMAP of co-embedded E6.5 cells, batch effects are observed between three studies and samples. **d,** UMAP of the same cells as in panel c with batch correction prior to integration (Stuart et al. 2019). The same UMAP is shown twice, colored by dataset (left, colors as in panel c) or cell type annotation (right). ExE: extraembryonic. EmVE: embryonic visceral endoderm. ExVE: extraembryonic visceral endoderm. **e**, UMAP visualization of co-embedding of data from *cells* at E8.5 generated on the 10x Genomics platform (Pijuan-Sala et al. 2019) and *nuclei* at E9.5 generated using sci-RNA-seq3 (Cao et al. 2019), after batch correction (Stuart et al. 2019). The same UMAP is shown twice, with colors highlighting cell states from either E8.5 (left) or E9.5 (right).

**Supplementary Figure 2.**
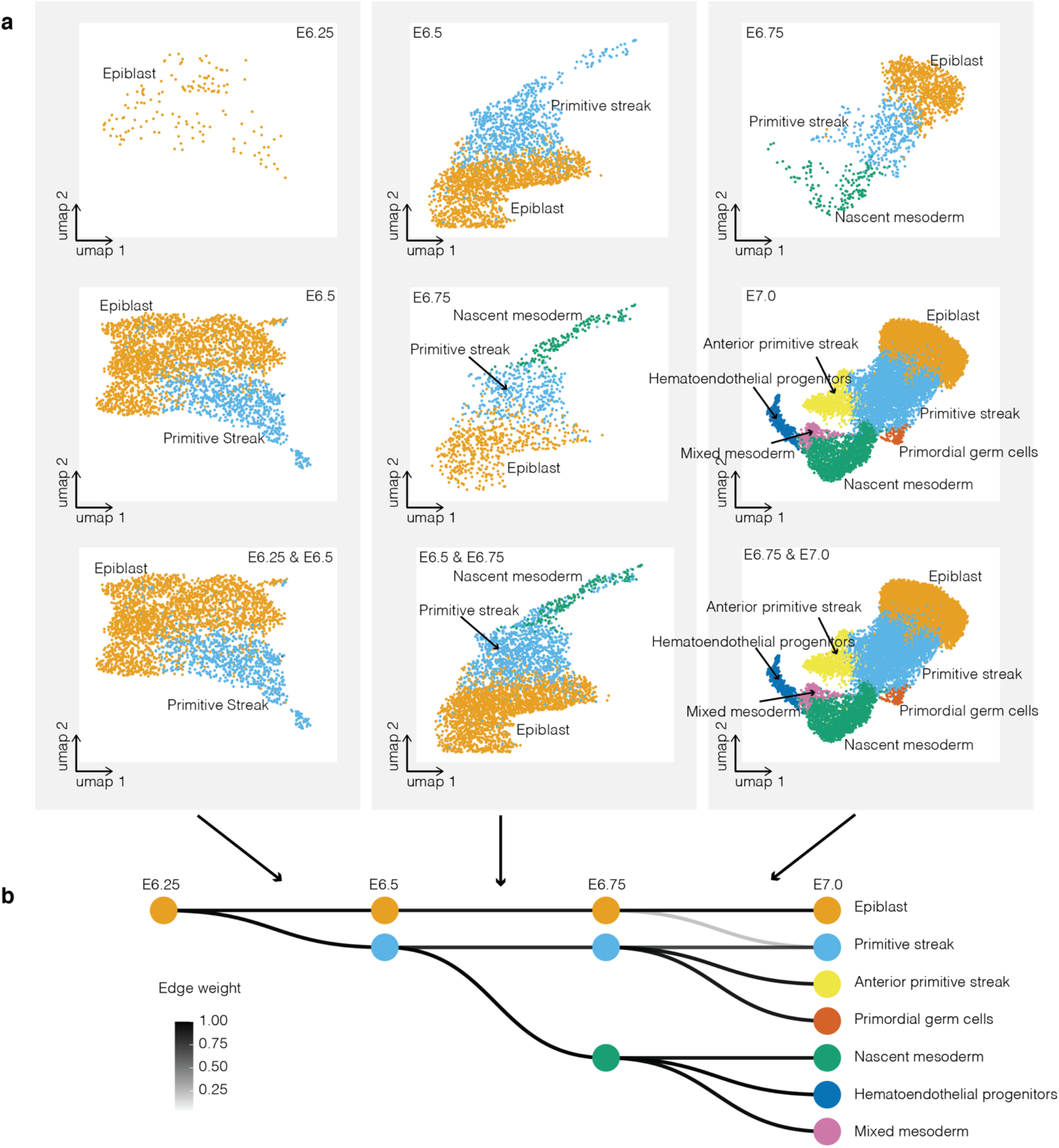
Inference of epiblast derivatives between E6.25 and E7.0. **a**, A portion of the UMAP corresponding to the epiblast and its inferred derivatives is shown for co-embeddings of E6.25 → E6.5 (left column), E6.5 → E6.75 (middle column) and E6.75 → E7.0 (right column). Within each column is the same UMAP visualization, but showing only cells from the earlier timepoint (top row), the later timepoint (middle row) or both timepoints (bottom row). **b**, Directed acyclic graph showing inferred relationships between cell states amongst early epiblast derivatives. All edges with weights above 0.2 are shown in grey scale.

**Supplementary Figure 3.**
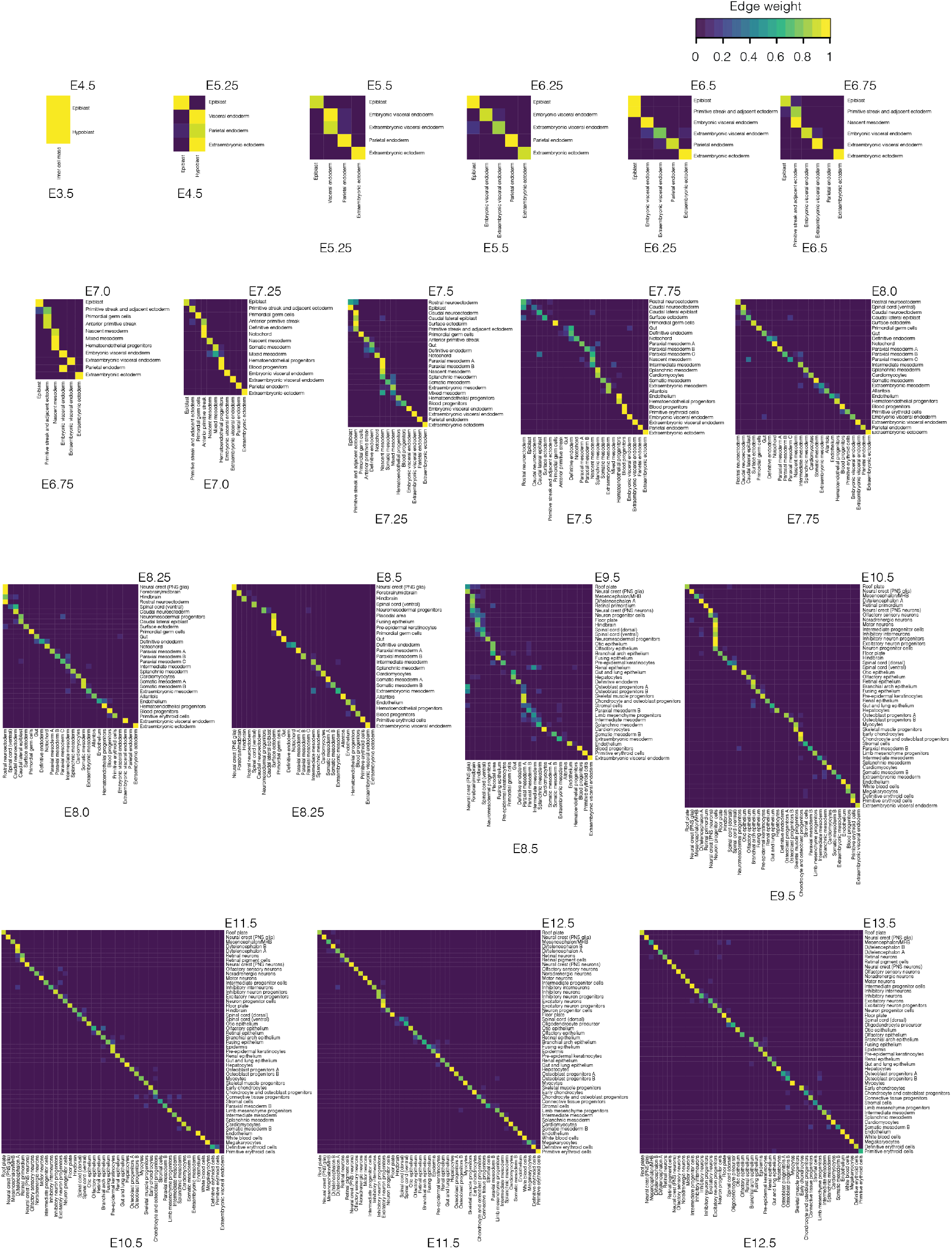
Heatmap of edge weights between cell states at each pair of adjacent timepoints. Each heatmap shows edge weights between all cell states at a given timepoint (rows) and potential pseudo-ancestral cell states from the immediately preceding timepoint (columns). Edge weights were calculated based on a *k*-nearest neighbor (*k*-NN) based heuristic that was applied to a co-embedding of separately annotated cells from the adjacent timepoints. The edge weights range from 0 to 1, and edges with weights greater than 0.2 were carried forward. PNS: peripheral nervous system. MHB: midbrain-hindbrain boundary.

**Supplementary Figure 4.**
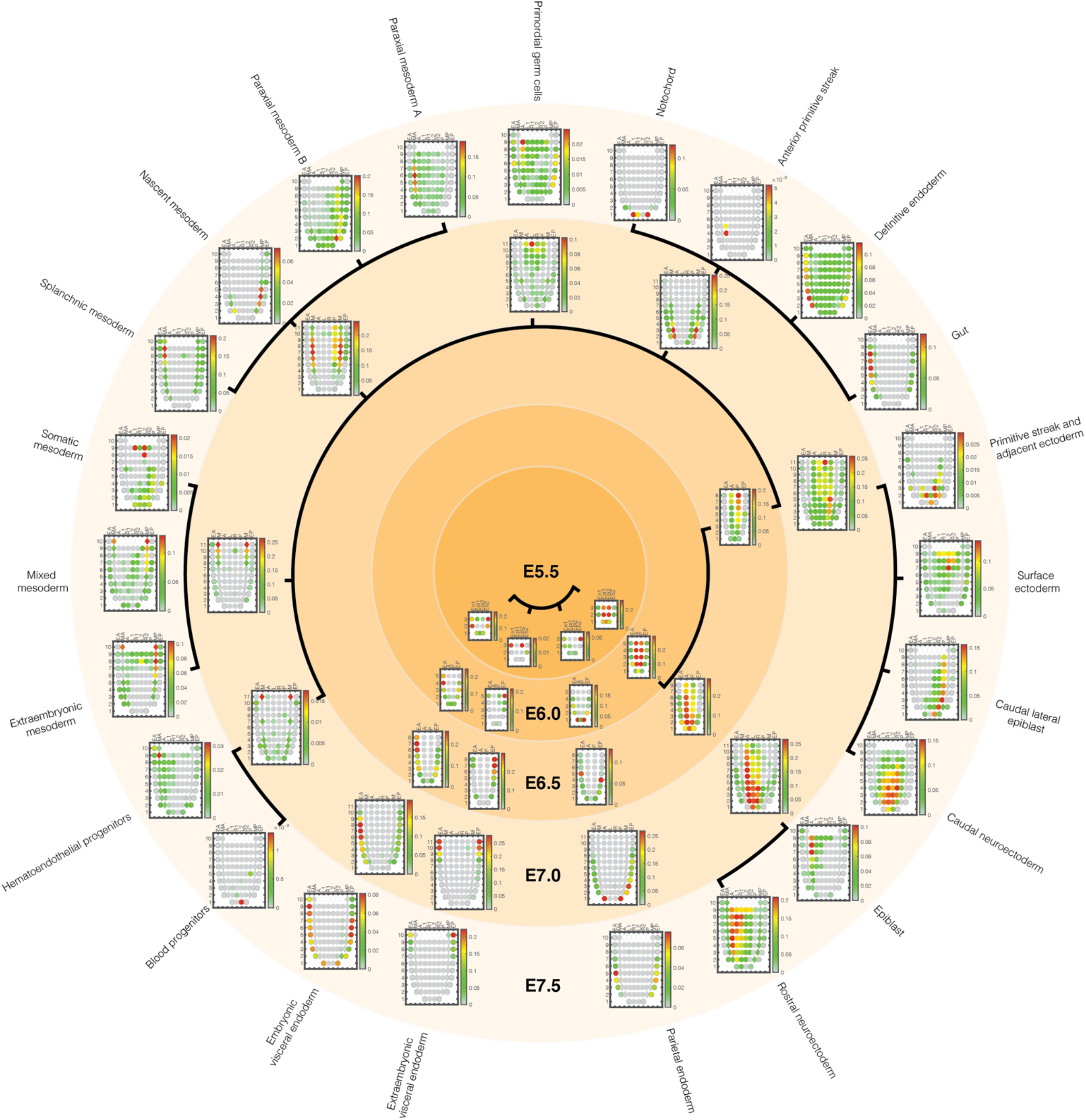
Estimated cell type proportions for different regions of the gastrulating mouse embryo, arranged by inferred cell type relationships over time. As described in Fig. 3a, inference of cell type contributor(s) to each spatial territory of the gastrulating mouse embryo based on the application of CIBERSORTx to GEO-seq data (Newman et al. 2019; Peng et al. 2019). Extraembryonic ectoderm excluded here because GEO-seq experiments only focused on the cell lineages derived from the inner cell mass (Peng et al. 2019). As scRNA-seq data from E6.0 was unavailable, we used data from E6.25 instead. Black edges correspond to edges between cell states over time estimated by TOME (only edges with the largest weights are shown). In each corn plot, each circle or diamond refers to a GEO-seq sample, and its weighted color to the estimated cell type composition. Corn plot nomenclature from (Peng et al. 2019). A, anterior; P, posterior; L, left lateral; R, right lateral; L1, anterior left lateral; R1, anterior right lateral; L2, posterior left lateral; R2, posterior right lateral; Epi1 and Epi2, divided epiblast; M, whole mesoderm; MA, anterior mesoderm; MP, posterior mesoderm; En1 and En2, divided endoderm; EA, anterior endoderm; EP, posterior endoderm.

**Supplementary Fig 5.**
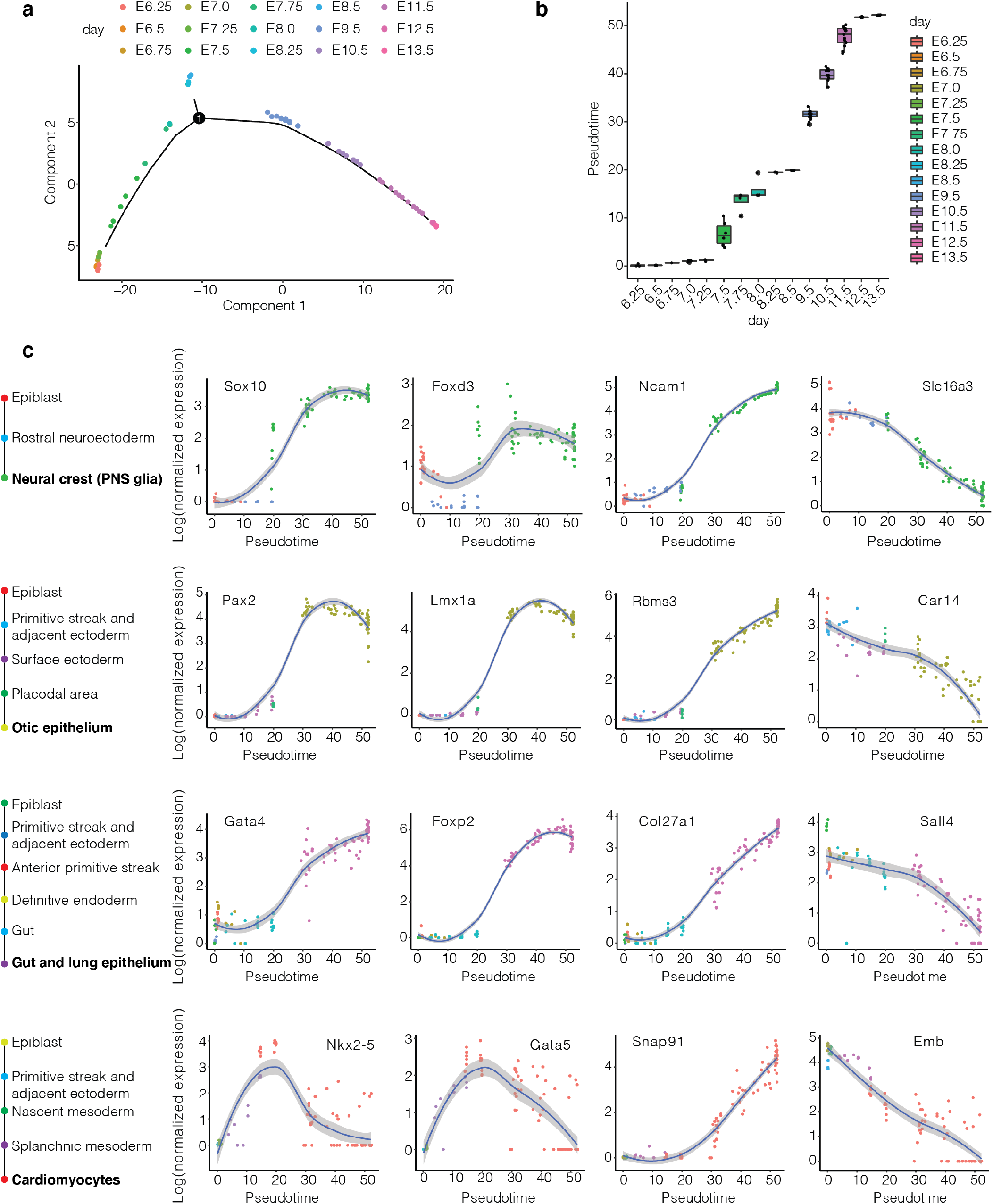
Inferring continuous molecular histories of individual cell types. **a**, Pseudotime trajectory analysis of pseudobulk RNA-seq profiles of mouse embryos. Briefly, epiblast-derived cells from individual embryos (or pools of embryos comprising each sample, in the case of (Pijuan-Sala et al. 2019)) were aggregated to create 99 pseudobulk samples, on which we performed pseudotime trajectory analysis. Each point in the resulting 2D embedding corresponds to an embryo, and the curve to pseudotime trajectory. **b**, Pseudotime of embryos from staged timepoints between E6.25 and E13.5. **c**, Smoothed expression profiles for four selected genes for each of four selected cell types (rows; one from each germ layer), along their inferred trajectories (key at left). We selected linear paths corresponding to strongest pseudo-ancestor edges, working back from each E13.5 cell state to the E6.25 epiblast cell state. The first and second columns of plots correspond to key regulators or marker genes, and the third and fourth columns to the genes most positively and negatively correlated with pseudotime, respectively. Each plotted point corresponds to gene expression within a cell state for an individual embryo. Pseudotime values (x-axes) as in panel b. Gene expression (y-axes) calculated as aggregated UMI within cell state normalized to total UMI per individual, followed by natural-log transformation. The inferred trajectory for the neural crest (PNS glia) spanned epiblast (E6.25 → E7.5), rostral neuroectoderm (E7.5 → E8.25), and neural crest (PNS glia) (E8.25 → E13.5). The inferred trajectory for the otic epithelium spanned epiblast (E6.25), primitive streak and adjacent ectoderm (E6.5 → E7.5), surface ectoderm (E7.5 → E8.25), placodal area (E8.5), and otic epithelium (E9.5 → E13.5). The inferred trajectory for the gut and lung epithelium spanned epiblast (E6.25), primitive streak and adjacent ectoderm (E6.5 → E6.75), anterior primitive streak (E7 → E7.25), definitive endoderm (E7.25 → E7.5), gut (E7.25 → E7.5), and gut and lung epithelium (E9.5 → E13.5). The inferred trajectory for the cardiomyocytes spanned epiblast (E6.25), primitive streak and adjacent ectoderm (E6.5), nascent mesoderm (E6.75 → E7.25), splanchnic mesoderm (E7.5 → E7.75), cardiomyocytes (E7.75 → E13.5). PNS: peripheral nervous system.

**Supplementary Figure 6.**
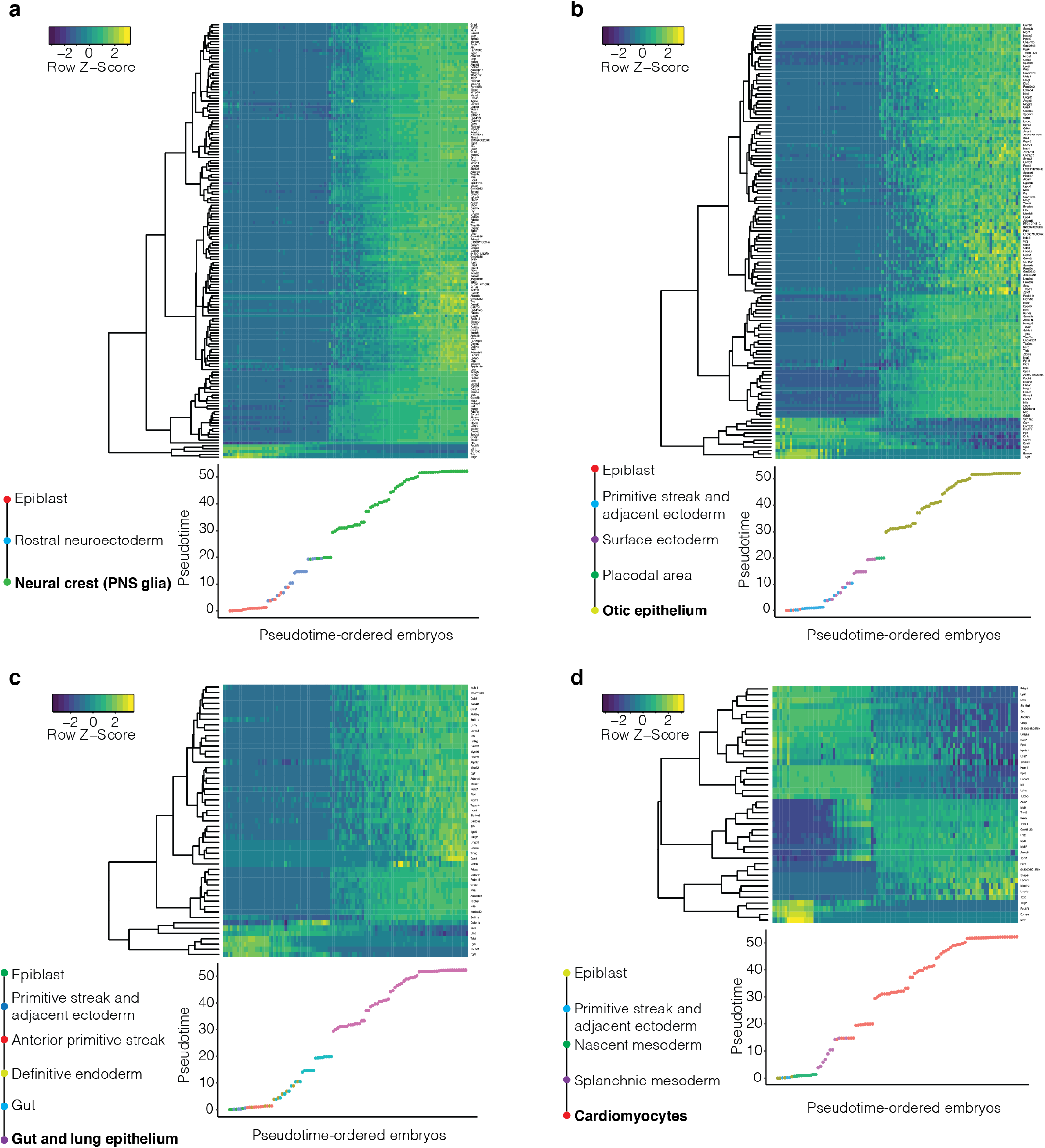
Gene dynamics across the inferred molecular trajectories of four selected cell types. **a**, 154 genes were identified as significantly associated with pseudotime of the neural crest (PNS glia) trajectory, based on linear regression with the origin of the data as a covariate. Genes with bonferroni-adjusted p-value (on the variable of pseudotime) < 0.05, absolute value of beta coefficient (on the variable of pseudotime) > 0.05, and absolute value of beta coefficient (on the variable of data identity) < 2 were retained and hierarchically clustered (y-axis of heatmap). The columns of the heatmap correspond to different embryos/samples, ordered by pseudotime (below) as shown in Supplementary Fig. 5a-b. **b**, 127 genes were identified as significantly associated with pseudotime of the otic epithelium trajectory. **c**, 51 genes were identified as significantly associated with pseudotime of the gut and lung epithelium trajectory. **d**, 42 genes were identified as significantly associated with pseudotime of the cardiomyocytes trajectory. Axes as well as thresholds for identifying genes and modules in panels b-d as in panel a. The inferred trajectory of each cell type included the same cell states as described in Supplementary Fig. 5c, in each case starting from epiblast. PNS: peripheral nervous system.

**Supplementary Figure 7.**
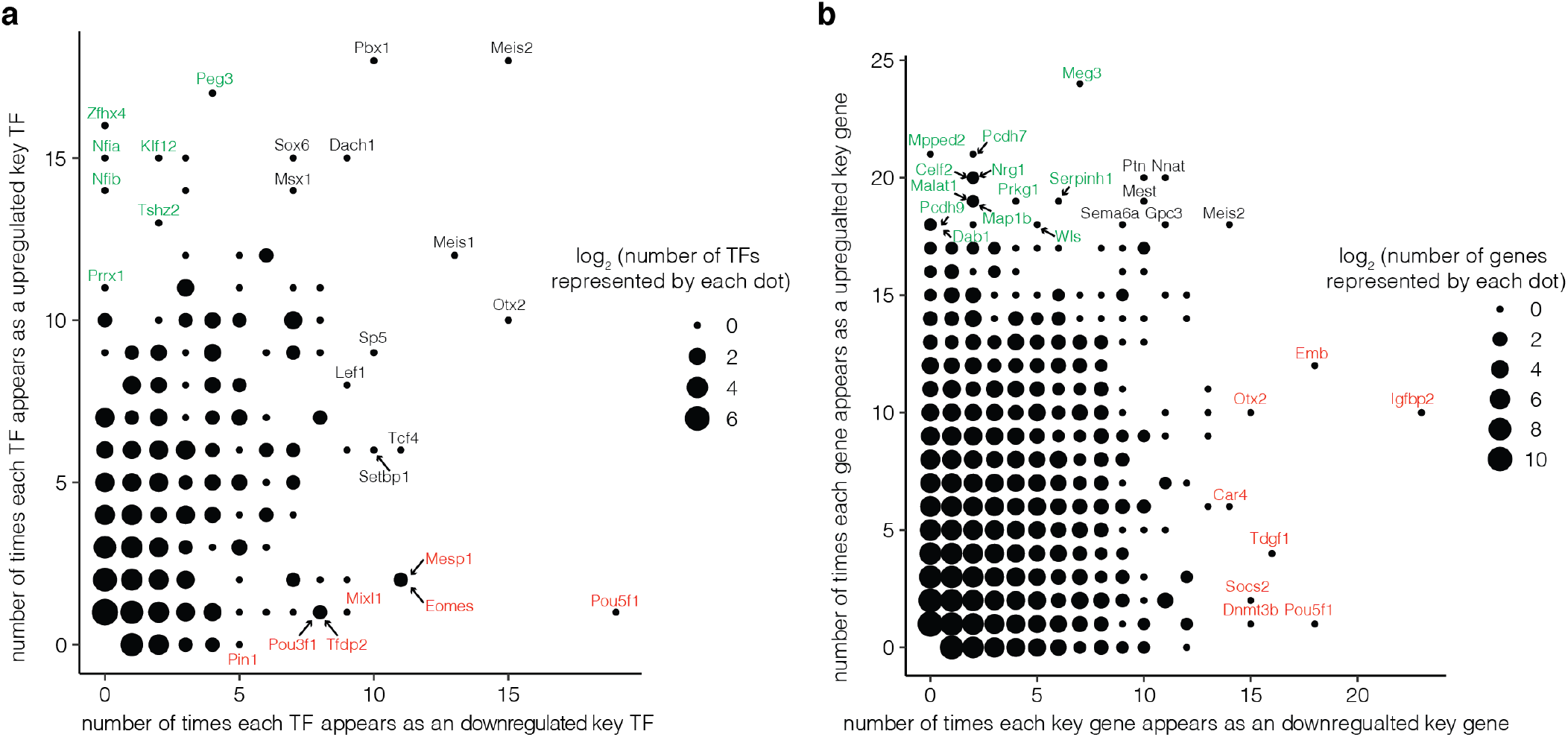
Recurrence of individual TFs or genes as candidate upregulated or downregulated key TFs or genes for mouse cell type specification. **a**, TFs that are most often nominated as downregulated key TFs, *e.g.* Pou5f1 (Oct4) are identified with red labels, while those most often nominated as upregulated key TFs, *e.g*. Zfhx4, are identified with green labels. Candidate key TFs frequently recurring in both sets are identified with black labels. The size of each dot corresponds to the number of TFs represented by it on a log2 scale. **b**, Genes that are most often nominated as downregulated key genes, *e.g.* Igfbp2, are identified with red labels, while those most often nominated as upregulated key genes, *e.g.* Meg3, are identified with green labels. Candidate key genes frequently recurring in both sets are identified with black labels. The size of each dot corresponds to the number of TFs represented by it on a log2 scale.

**Supplementary Figure 8.**
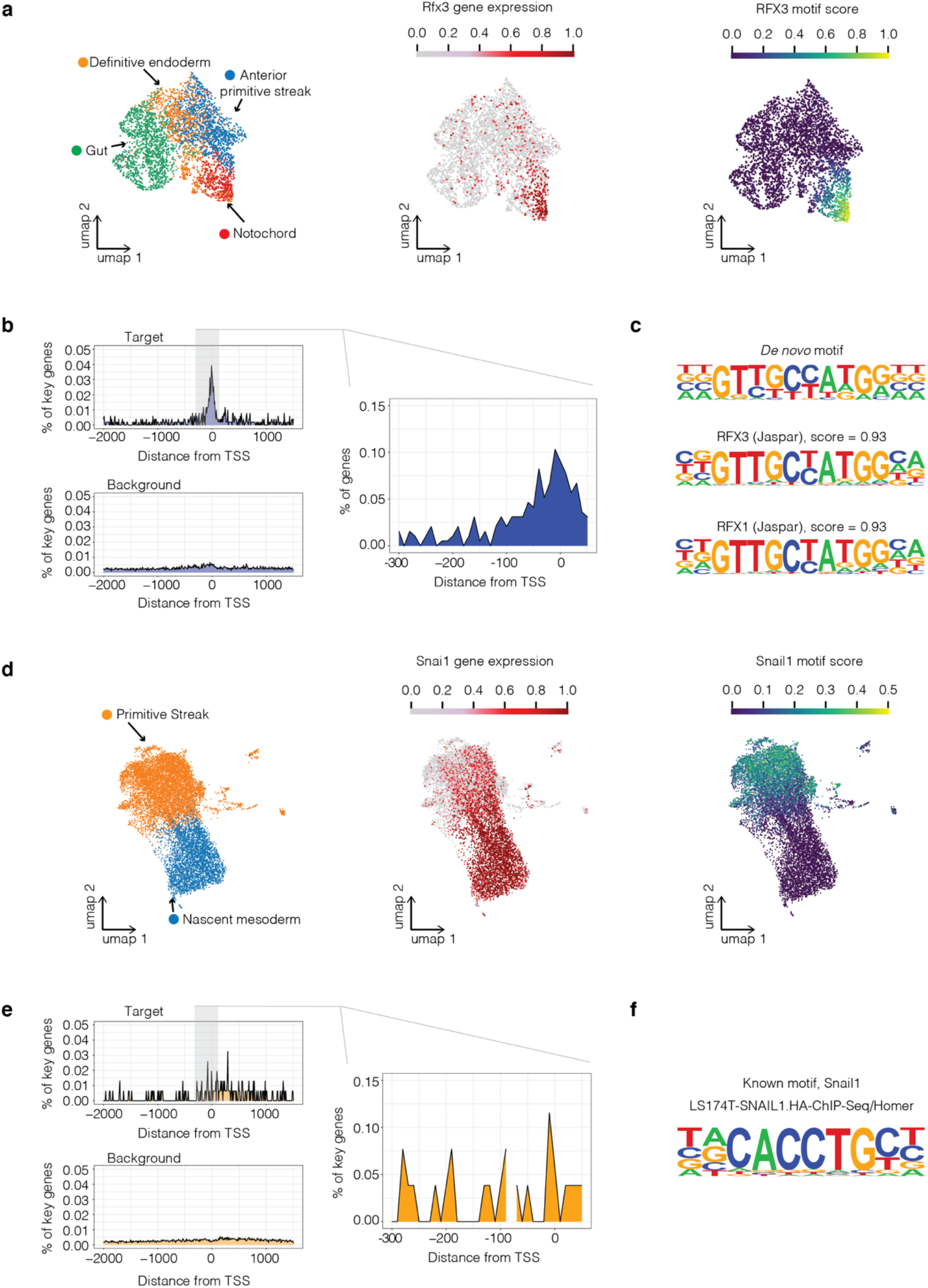
Correlation between key TF expression and up- or down-regulation of putative targets of regulation. **a**, UMAP visualization of co-embedded cells from cell states including anterior primitive streak, definitive endoderm, gut, and notochord (mouse E7.25 → E7.5) colored by cell type (left), Rfx3 gene expression (middle) or RFX3 motif score (right), respectively. The RFX3 motif score for each cell was calculated by averaging the gene expression of 135 key genes for notochord emergence bearing this motif in their core promoters, and then subtracting the mean expression of a reference set of randomly sampled genes, using the score_genes function of Scanpy (Wolf, Angerer, and Theis 2018). **b**, Positional bias of RFX3 binding motif along the core promoters of key genes for notochord emergence (right panel), an expanded region for key genes for notochord emergence (left top panel), or an expanded region for background (left bottom panel). The y-axes indicate the % of key genes or background genes with the RFX3 motif with 10 bp bins. **c**, The motif logo of the top de novo identified motif for notochord emergence and its two best alignments in the known motif database. **d**, UMAP visualization of co-embedded cells from cell states including primitive streak and nascent mesoderm (mouse E6.5 → E7.25) colored by cell types (left), Snai1 gene expression (middle) or SNAIL1 motif score (right), respectively. The SNAIL1 motif score was calculated as in panel a, based on 21 key genes for nascent mesoderm emergence bearing this motif in their core promoters. **e**, Positional bias of SNAI1 binding motif along the core promoters of key genes for nascent mesoderm emergence (right panel), an expanded region for key genes for nascent mesoderm emergence (left top panel), or an expanded region for background (left bottom panel). The y-axes indicate the % of key genes or background genes with the SNAIL1 motif with 10 bp bins. **f**, The known motif logo of SNAIL1.

**Supplementary Fig 9.**
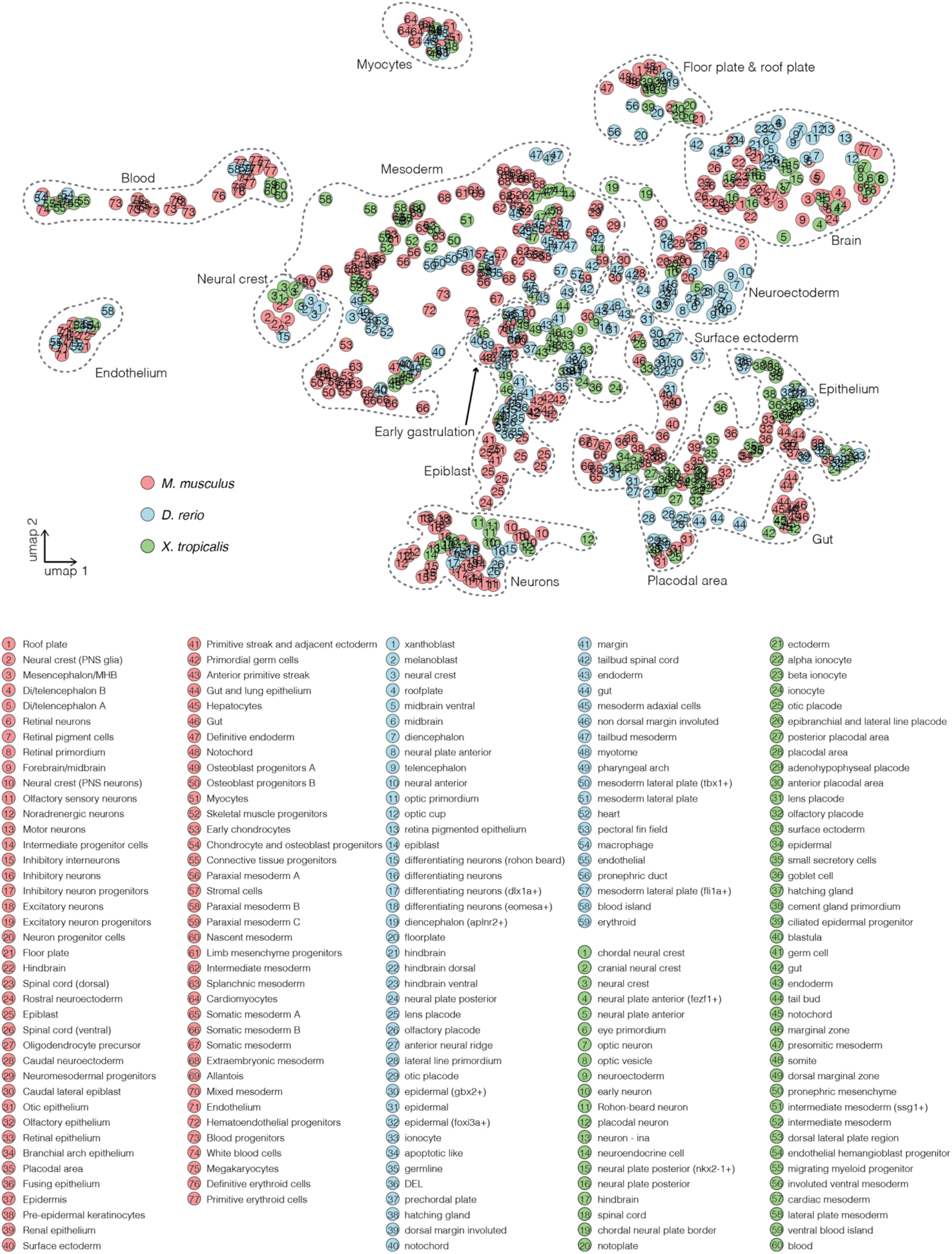
Co-embedding of 765 cell states from three species by integrating their transcriptional features. For cell states spanning multiple timepoints, cells from each timepoint were treated separately for the purposes of this analysis. To create the transcriptional feature of each cell state (*i.e.* a pseudo-cell), we first averaged cell-state-specific UMI counts, normalized by the total count, multiplied by 100,000 and natural-log-transformed after adding a pseudocount. We then divided all resulting 765 pseudo-cells from the three species into four groups: the mouse single-cell group (n = 142), the mouse single-nucleus group (n = 226), the zebrafish group (n = 205), and the frog group (n = 192), and performed the anchor-based batch correction (Stuart et al. 2019). UMAP visualization shows co-embedded pseudo-cells from the mouse (red), the zebrafish (blue), and the frog (green). Each circle corresponds to a pseudo-cell, and the numbers correspond to the cell state labels shown below. The grey dotted curves (manually added) highlight 15 major groups, each including representatives from all three species. Cell states from the extraembryonic lineages (Inner cell mass, hypoblast, parietal endoderm, extraembryonic ectoderm, visceral endoderm, embryonic visceral endoderm, and extraembryonic visceral endoderm for the mouse; blastomere, EVL, periderm, forerunner cells for the zebrafish) were excluded from this analysis. For E6.5 of mice, we only used cells from a single study (Pijuan-Sala et al. 2019). PNS: peripheral nervous system. MHB: midbrain-hindbrain boundary. DEL: deep cell layer. EVL: enveloping layer.

**Supplementary Figure 10.**
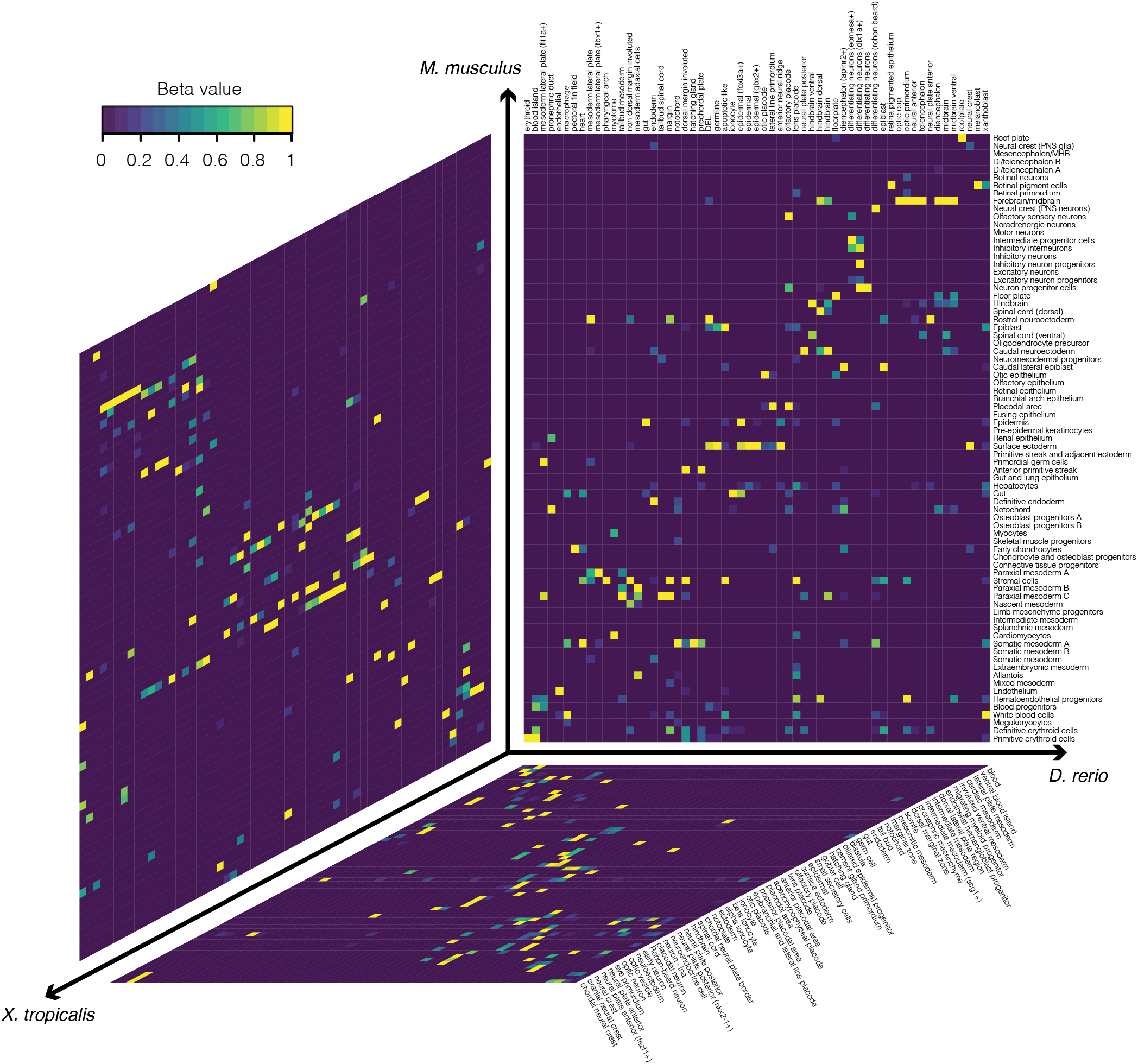
Correlated cell types between species based on non-negative least-squares (*NNLS*) regression. To identify correlated cell types between each pair of species, we averaged cell-type-specific UMI counts, normalized by the total count, multiplied by 100,000 and natural-log-transformed after adding a pseudocount. Extraembryonic cell types (inner cell mass, hypoblast, parietal endoderm, extraembryonic ectoderm, visceral endoderm, embryonic visceral endoderm, and extraembryonic visceral endoderm for the mouse; blastomere, EVL, periderm, forerunner cells for the zebrafish) were excluded from this analysis. For mouse E6.5, we only used cells from a single study (Pijuan-Sala et al. 2019). We then applied *NNLS* regression to predict the gene expression of target cell type (*T_a_*) in dataset A with the gene expression of all cell types (*M_b_*) in dataset B: *T_a_* = *β*_0a_ + *β*_1a_*M_b_*, based on the union of the 1,200 most highly expressed genes and 1,200 most highly specific genes in the target cell type. We then switched the roles of datasets A and B, *i.e.* predicting the gene expression of target cell type (*T_b_*) in dataset B from the gene expression of all cell types (*M_a_*) in dataset A: *T_b_* = *β*_0b_ + *β*_1b_*M_a_*. Finally, for each cell type a in dataset A and each cell type b in dataset B, we combined the two correlation coefficients: *β* = 2(*β*_ab_ + 0.001)(*β*_ba_ + 0.001), a statistic for which high values reflect reciprocal, specific predictivity. The combined *β* value was normalized by dividing by the maximum value of each column (zf for mm vs. zf, xp for mm vs. xp, and xp for zf vs. xp). Shown here is a heat map of the normalized β values between 77 cell types from the mouse, 59 cell types from the zebrafish, and 60 cell types from the frog. The order of cell types listed in the heat map is the same as each cellular trajectory plot (Fig. 1c; Fig. 5b**-c**). PNS: peripheral nervous system. MHB: midbrain-hindbrain boundary. DEL: deep cell layer.

**Supplementary Figure 11.**
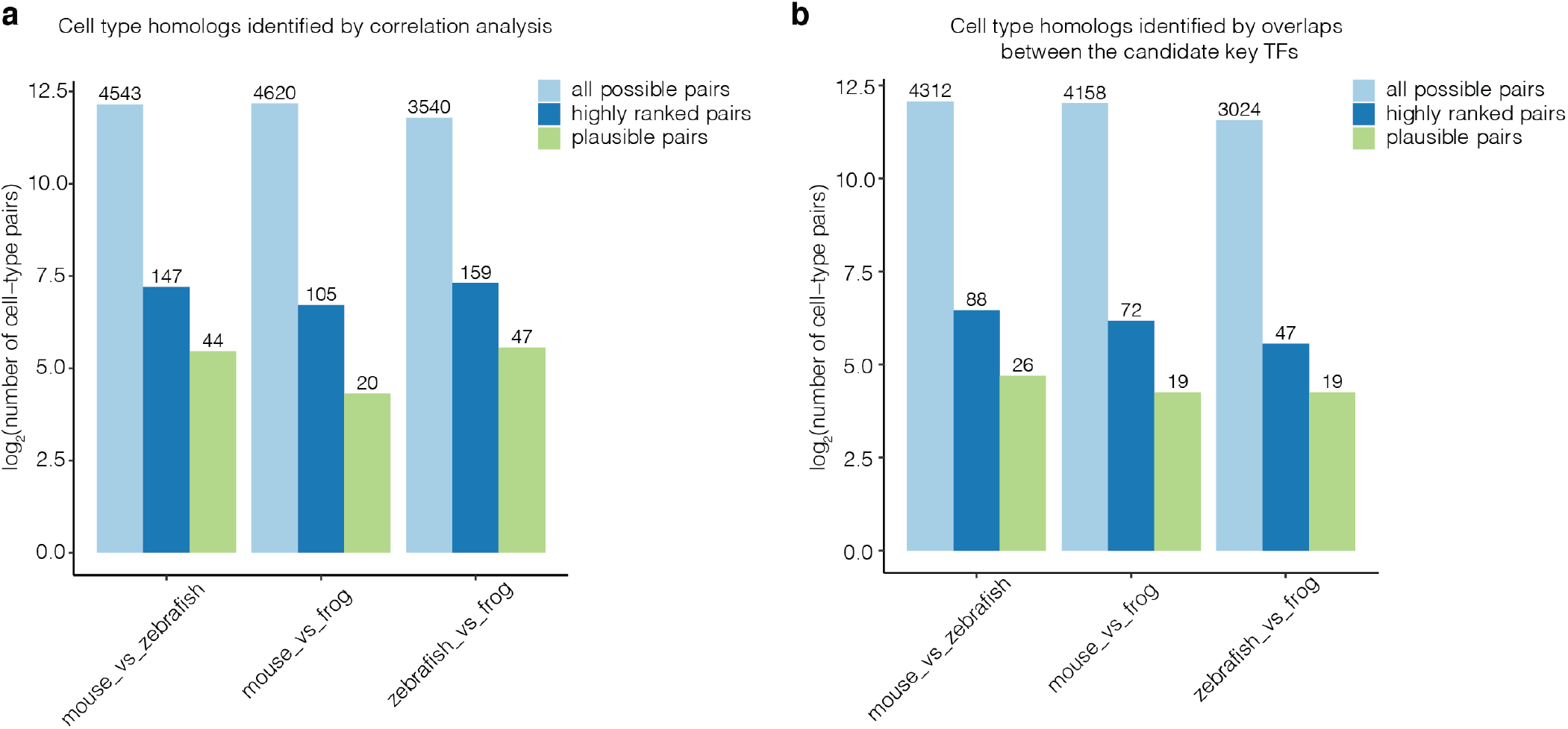
The log-scaled number of all possible pairs, highly ranked pairs, and biologically plausible pairs of cell types identified in pairwise comparisons of three species (mouse, zebrafish, frog) by two strategies. **a**, The log2-scaled number of all possible pairs, highly ranked pairs, and biologically plausible pairs of cell types evaluated by non-negative least-squared (*NNLS*) regression. “All possible pairs” refers to all potential cell type pairings considered; “highly ranked pairs’’ refer to pairings with *β* > 1e-4 and that ranked highly from the perspective of both species; “plausible pairs” refer to pairings which were retained after manual review for biological plausibility (**Supplementary Table 21**). Actual numbers shown above each bar, with y-axis on log2-scale. **b**, The log2-scaled number of all possible pairs, highly ranked pairs, and biologically plausible pairs of cell types evaluated on the basis of overlapping, orthologous candidate key TFs. “All possible pairs” refers to all potential cell type pairings considered; “highly ranked pairs’’ refer to pairings with estimated relative likelihoods more extreme than 99% of permutations; “plausible pairs” refer to pairings which were retained after manual review for biological plausibility (**Supplementary Table 22**). Actual numbers shown above each bar, with y-axis on log2-scale.

